# Cryo-EM structure of MukBEF reveals DNA loop entrapment at chromosomal unloading sites

**DOI:** 10.1101/2021.06.29.450292

**Authors:** Frank Bürmann, Louise F.H. Funke, Jason W. Chin, Jan Löwe

## Abstract

The ring-like structural maintenance of chromosomes (SMC) complex MukBEF folds the genome of *Escherichia coli* and related bacteria into large loops, presumably by active DNA loop extrusion. MukBEF activity within the replication terminus macrodomain is suppressed by the sequence specific unloader MatP. Here we present the complete atomic structure of MukBEF in complex with MatP and DNA as determined by electron cryomicroscopy (cryo-EM). The complex binds two distinct DNA double helices corresponding to the arms of a plectonemic loop. MatP-bound DNA threads through the MukBEF ring, while the second DNA is clamped by the kleisin MukF, MukE and the MukB ATPase heads. Combinatorial cysteine cross-linking confirms this topology of DNA loop entrapment *in vivo*. Our findings illuminate how a class of near-ubiquitous DNA organizers with important roles in genome maintenance interacts with the bacterial chromosome.

**Highlights:**

- Complete atomic structures of the bacterial SMC complex MukBEF on and off DNA.
- MukBEF entraps two DNA double helices when bound to the unloader MatP.
- *In vivo* topology of DNA loop entrapment determined by cysteine cross-linking.
- Arms of the DNA loop thread through separate compartments of MukBEF.

**Graphical abstract:** 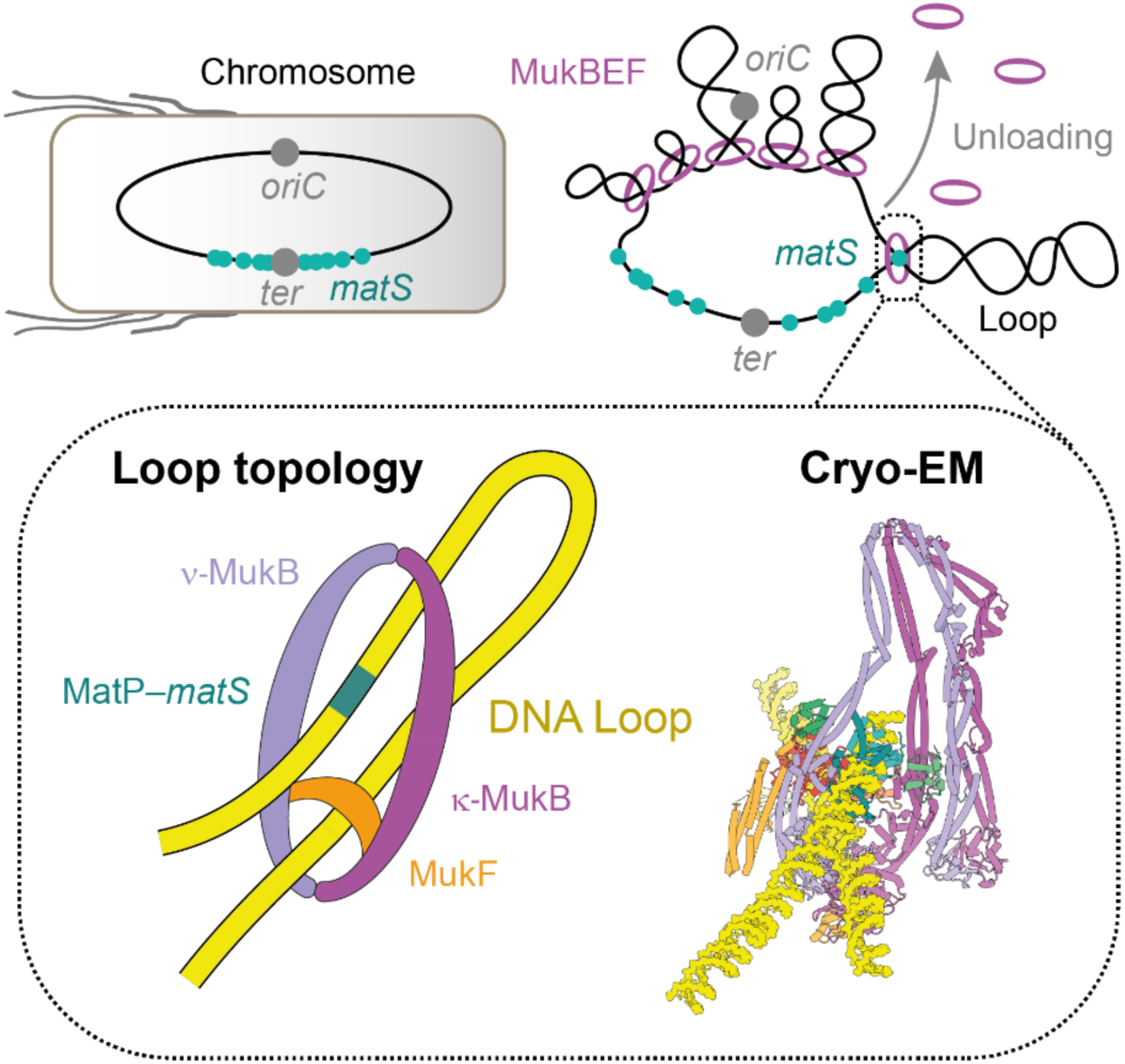

## Introduction

Associations between molecules due to their topology are known as mechanical bonds (Stoddart, 2009). In eukaryotes as well as prokaryotes, ring-like SMC complexes are thought to structure chromosomes via mechanical bonds with DNA (also referred to as ‘DNA entrapment’), and active DNA loop extrusion (Davidson and Peters, 2021; Hassler et al., 2018; Mäkelä and Sherratt, 2020a; Nasmyth, 2001; Yatskevich et al., 2019). These activities have been suggested to enable or facilitate processes such as lengthwise condensation of chromosomes, sister chromatid cohesion, regulation of interactions between enhancers and distant promoters, disentangling of replicated DNA by topoisomerases, DNA recombination, and DNA double-strand break repair. Support for DNA entrapment, whereby the SMC complex fully encircles the nucleic acid polymer, comes from experiments probing DNA association after high salt treatment or, more stringently, chemical circularization and denaturation of the complex (Cuylen et al., 2011; Haering et al., 2008; Ivanov and Nasmyth, 2005; Kanno et al., 2015; Murayama and Uhlmann, 2013; Niki and Yano, 2016; Wilhelm et al., 2015). How DNA entrapment is achieved on the molecular level is less well understood, as SMC complexes contain multiple topological compartments which can or could accommodate one or more DNA double strands (Chapard et al., 2019; Collier et al., 2020; Higashi et al., 2020; Shi et al., 2020; Vazquez Nunez et al., 2019).

At the core of SMC complexes, such as cohesin, condensin, Smc5–6, Smc–ScpAB, and MukBEF, is a tripartite ring composed of two SMC proteins and a kleisin. SMC proteins contain a 50- nm-long intra-molecular anti-parallel coiled-coil ‘arm’, which can fold over at an ‘elbow’ in MukBEF, cohesin and condensin (Bürmann et al., 2019; Lee et al., 2020; Niki et al., 1992). The arm separates a ‘hinge’ dimerization domain from an ABC-type ATPase ‘head’ domain, which undergoes cycles of ATP-dependent dimerization (called ‘engagement’), ATP hydrolysis, and disengagement (Hopfner et al., 2000). The kleisin bridges hinge-dimerized SMC proteins in an asymmetric arrangement, whereby its N- and C-terminal domain bind the SMC ‘neck’ and ‘cap’ surfaces, respectively (Bürmann et al., 2013; Gligoris et al., 2014; Haering and Gruber, 2016; Haering et al., 2004; Woo et al., 2009; Zawadzka et al., 2018). The neck is located at the very head-proximal region of the arm, whereas the cap is part of the head domain. The neck-bound SMC is designated *ν*-SMC (ny for neck), and the cap-bound subunit is designated *κ*- SMC (kappa for cap).

MukBEF is the SMC complex of *E. coli* and other enterobacteria. Its SMC subunit is MukB, which associates with the kleisin MukF (Woo et al., 2009; Zawadzka et al., 2018). MukF binds the dimeric KITE protein MukE, which is structurally related to ScpB of prokaryotic Smc–ScpAB and Nse1–3 of the Smc5–6 complex (Palecek and Gruber, 2015). MukB_2_E_2_F assemblies (hereafter ‘MukBEF monomers’) dimerize via MukF (Fennell-Fezzie et al., 2005; Woo et al., 2009) into MukB_4_E_4_F_2_ complexes (also called ‘MukB dimers of dimers’, hereafter simply ‘MukBEF dimers’), which are the functional form (Badrinarayanan et al., 2012; Rajasekar et al., 2019).

MukBEF organizes large fractions of the *E. coli* chromosome and is essential for chromosome segregation and cell survival under conditions of fast growth (Danilova et al., 2007; Hiraga et al., 1989; Lioy et al., 2018). In the replication terminus macrodomain (*Ter*), MukBEF activity is suppressed by unloading at *matS* sites, which are the signature sequences of *Ter* (Lioy et al., 2018; Mäkelä and Sherratt, 2020b; Mercier et al., 2008; Nolivos et al., 2016). Unloading of MukBEF drastically changes the loop-size distribution of *Ter* compared to other chromosomal macrodomains, a process which depends on the *matS*-binding protein MatP. Although MukBEF function relies on the full ATPase cycle (Badrinarayanan et al., 2012; Woo et al., 2009), association with *matS* requires ATP-dependent head engagement only (Nolivos et al., 2016). Head engagement and ATP hydrolysis have been suggested to mediate unloading of cohesin in eukaryotes, which involves dissociation of the neck/kleisin interface (Chan et al., 2012; Muir et al., 2020; Murayama and Uhlmann, 2015). This interface also disengages during the ATPase cycle of condensin, raising the possibility that SMC complexes may use related mechanisms for DNA unloading (Hassler et al., 2019).

Here, using cryo-EM single particle analysis we discovered that MukBEF entraps two distinct DNA double helices when bound to the unloader MatP. The DNAs reside in separate compartments, which are located inside the large circumference of the tripartite ring and in a much smaller clamp at the ATPase heads, respectively. Topological mapping by chemical circularization of endogenous MukBEF in cells suggests that these compartments enclose each arm of a DNA loop *in vivo*. Our findings illuminate how MukBEF can entrap DNA loops, and how these loops are primed for unloading from the complex.

## Results

### MukBEF–MatP entraps two DNA double helices in topologically separate compartments

Initial attempts to determine the structure of *E. coli* MukBEF were unsuccessful. We then recombinantly produced MukBEF complexes from 11 different species in *E. coli* and identified the complex from *Photorhabdus thracensis* as a suitable candidate for structure determination by cryo-EM. The complex has 78 % sequence identity to its *E. coli* homologue and stably co-purified with the *E. coli* acyl-carrier protein AcpP, an essential protein that is a binding partner of *E. coli* MukBEF (Niki et al., 1992; Prince et al., 2021). *E. coli* AcpP is 85 % identical to *P. thracensis* AcpP. The purified complex had an estimated stoichiometry of MukB_2_E_4_F_2_–AcpP_2_, which is a MukB_2_–AcpP_2_ unit short of the MukB_4_E_4_F_2_ dimer expected from live-cell single-molecule microscopy (Badrinarayanan et al., 2012). We reasoned that this was either due to dissociation of MukBEF dimers during purification, or due to overproduction of MukEF leading to saturation and incomplete assembly of the complex. We therefore titrated the preparation with MukB_2_–AcpP_2_ to reconstitute intact MukB_4_E_4_F_2_ dimers. This almost quantitatively shifted MukBEF to a smaller elution volume in size exclusion chromatography (SEC), indicating the formation of the physiological complex (**Figure 1A**).

**Figure 1.**
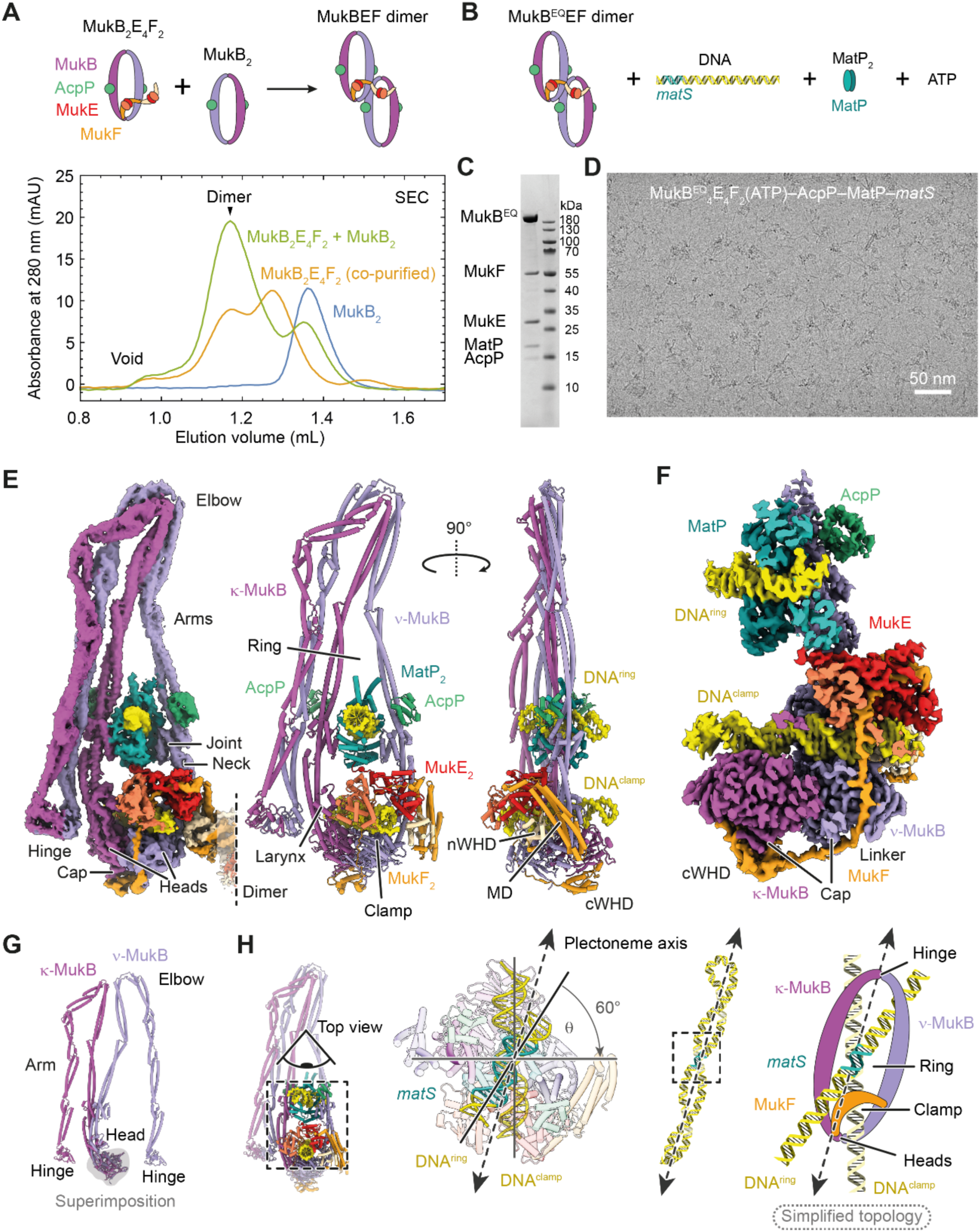
Cryo-EM structure of MukBEF–MatP bound to two distinct DNA double helices. (**A**) Reconstitution of MukBEF dimers. Co-purified MukBEF and free MukB were mixed (top) and resolved by SEC (bottom). (**B**) Composition of the MukBEF–MatP–*matS* sample used for structure determination. (**C**) Coomassie stained SDS-PAGE gel of the reconstituted complex used for cryo-EM. (**D**) Example micrograph of the sample used for structure determination. (**E**) 4.6 Å resolution cryo-EM density map (left) and complete atomic model (middle, right) of the DNA bound MukBEF–MatP monomer. (**F**) Slice through a 3.1 Å resolution cryo-EM density map of the DNA binding region of MukBEF–MatP. (**G**) *κ*-MukB and *ν*-MukB superimposed on the head domain. The arms adopt radically different conformations. (**H**) DNA binding topology on plectonemic loops inferred from the DNA crossing angle. The schematic on the right shows the simplified topology used for clarity throughout, with the in-reality elbow folded conformation flattened into a ring. See also **Figures S1**, **S2** and **S3**.

To gain insights into how MukBEF interacts with *matS* sites during chromosomal unloading, we purified *P. thracensis* MatP, identified a cognate high-affinity *matS* site (**Figure S1**) and introduced the E1407Q mutation (hereafter MukB^EQ^) into MukB to slow down ATP hydrolysis (Woo et al., 2009). We then reconstituted a complex between MukB^EQ^EF dimers, MatP, and an 80 bp DNA oligonucleotide containing *matS* close to one end, in the presence of ATP and magnesium ions (**Figures 1B** and **1C**). The sample was then imaged by cryo-EM in vitreous ice (**Figure 1D**).

We obtained a reconstruction of the DNA-bound MukBEF monomer part at an overall nominal resolution of 4.6 Å (**Figures 1E** and **S2**). Focussed classification and refinement produced a map of 3.1 Å resolution for the more rigid head module, with clearly resolved ATP and magnesium ions mediating head engagement (**Figure 1F**, **S3A**). This allowed the construction of a complete atomic model for the complex, facilitated by previous crystallographic information for individual parts (PDB: 3EUH, 3EUJ, 3IBP, 3VEA, 6H2X) (Bürmann et al., 2019; Dupaigne et al., 2012; Kreamer et al., 2018; Li et al., 2010; Woo et al., 2009).

MukBEF adopts a compact and highly asymmetric conformation, with its ATPase heads bridged by the kleisin MukF (**Figure 1E**). The heads and hinge of MukB are brought into proximity by folding at the elbow. In addition to the elbow and the ‘joint’ at the heads (Diebold-Durand et al., 2017), the MukB arms contain several other coiled-coil discontinuities (Weitzel et al., 2011). Plasticity in these regions allows *κ*-MukB and *ν*-MukB to adopt radically different conformations and thus break homodimer symmetry (**Figure 1G**).

The head-proximal arms of MukB are opened to allow accommodation of two distinct DNA double helices (**Figures 1E-1F, 1H** and **2A**). One DNA is bound by MatP and threads through the inter-arm space near the joint. The other is clamped by the MukB heads, MukF, and MukE. The chain of the kleisin MukF is resolved between residues 22-440, which represents 95 % of the protein (**Figures 1F** and **2B**). It partitions the two DNAs into topologically separated compartments, the ‘ring’ delimited by MukF and the MukB arms, and the ‘clamp’ delimited by MukF and the MukB heads (**Figures 1E** and **1H**). The DNA double helices have a crossing angle of 60°, which is close to what has been estimated for negatively supercoiled plectonemes (Rawdon et al., 2016). This suggests that in the context of an intact chromosome they may originate from a single plectonemic loop, with MukBEF binding across the long axis of the loop (**Figure 1H**). This hypothesis will be explored below.

**Figure 2.**
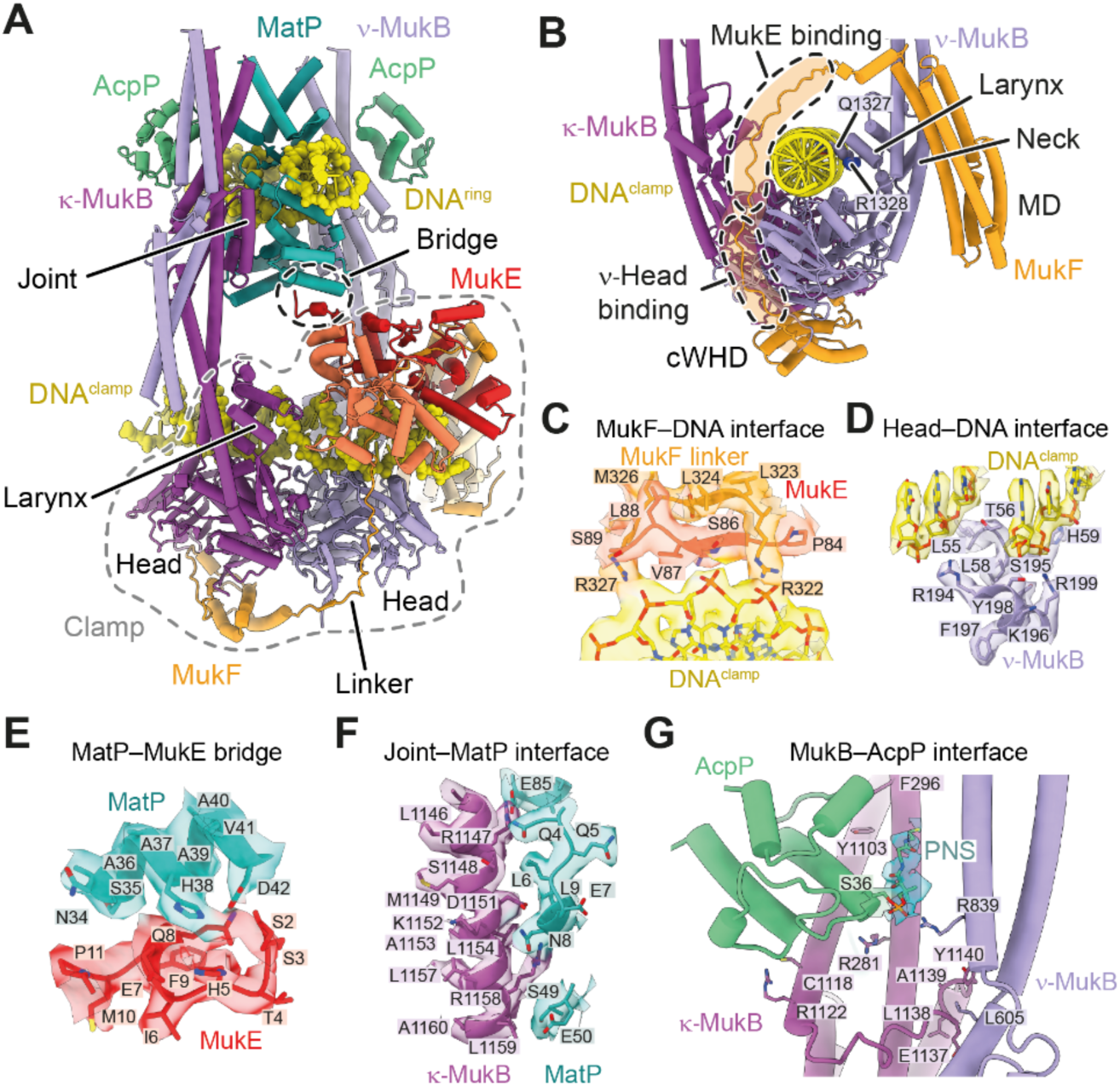
DNA binding and subunit interfaces of the MukBEF–MatP complex. (**A**) Model of the DNA-bound head module. (**B**) Path of the kleisin MukF and DNA contacts of the MukB larynx. (**C**) Interface between the MukF linker and the clamped DNA. (**D**) Interface between the top surface of the *ν*-MukB head and the clamped DNA. (**E**) Interface between MukE and MatP. (**F**) Interface between the MukB joint and MatP. (**G**) Interfaces between MukB and AcpP, and between the *κ*-MukB joint and the hinge-proximal arm of *ν*-MukB. See also Figures S2 and S3.

### The clamp contacts DNA across all core subunits

The clamp has a highly asymmetric architecture imposed by the kleisin MukF. MukF is comprised of a C-terminal winged-helix domain (cWHD), a four-helix bundle forming the middle domain (MD), and an N-terminal winged-helix domain (nWHD). The cWHD and MD are connected by a 64 amino acid linker, which contains the MukE binding sites (**Figure 2B**). The cWHD binds the cap of its cognate *κ*-SMC (**Figure 1F**), similar to the corresponding interface in other SMC complexes (Bürmann et al., 2013; Haering et al., 2004; Hassler et al., 2019; Woo et al., 2009). The MD which can bind the MukB neck (Zawadzka et al., 2018) is in a position roughly equivalent to binding sites between the kleisins and *ν*-SMCs of cohesin, condensin and Smc-ScpAB (Bürmann et al., 2013; Gligoris et al., 2014; Hassler et al., 2019) (**Figure S3B**). The MD is, however, structurally unrelated to the N-terminal alpha-helical domain (nHD) of these kleisins.

MukB forms a homodimer, thus both MukB subunits contain MukF binding sites at cap and neck. However, the *ν*-MukB does not form the cap/cWHD interface and *κ*-MukB does not associate with an MD. This asymmetric configuration is enabled by two separate steric occlusion mechanisms. At the *ν*-MukB cap, binding of the MukF linker to *ν*-MukB prevents binding of a cWHD, as has been observed before (Woo et al., 2009) (**Figures 1F** and **2B**). In addition, the neck of *κ*-MukB is occluded by the hinge (**Figure 1E**), which prevents binding of a MD. These mechanisms preclude recruitment of additional MukEF subunits to the complex and thus prevent chaining of MukBEF monomers into higher order polymers.

Within the clamp, all core subunits of MukBEF are in contact with DNA. The MukF linker is guided over the clamped DNA by the MukE dimer, which also binds DNA along its central cleft (**Figures 2A** and **2B**). The linker itself contacts the phosphate backbone with R322 and R327 (**Figure 2C**). The MukB heads bind the clamped DNA along their top surface (**Figure 2D**). The ‘larynx’ of *ν*-MukB provides additional DNA contacts with Q1327 and R1328 (**Figure 2B**). This globular domain is situated at the base of the neck and is not present in most other SMC proteins (**Figure S3C**). Interestingly, the nHD of cohesin’s kleisin Rad21 provides DNA contacts that are located in a position similar to the larynx (**Figure S3B**).

The overall architecture of the MukBEF clamp appears analogous to what has been observed for the nuclease clamp in SbcCD (Rad50–Mre11) and the HAWK clamp in cohesin (Collier et al., 2020; Higashi et al., 2020; Käshammer et al., 2019; Shi et al., 2020) (**Figure S3D**). The KITE MukE is unrelated to any subunit in these complexes, however, other KITE-based SMC complexes, such as Smc5–6 and Smc–ScpAB, may clamp DNA in a manner similar to MukBEF. We conclude that DNA binding on top of the ATPase heads is a common principle between different SMC complexes, whereas the non-SMC subunits create structurally divergent but topologically equivalent clamps.

### MatP and MukE bridge the two DNAs

Whereas the clamped DNA is contacted by all core subunits of the complex, the DNA inside the ring is mostly bound by MatP, with only K1178 in the MukB joint contacting the phosphate backbone. The MatP dimer recognizes *matS* by inserting its *α*4 and *β*1 elements into the major grove, as has been determined for MatP–*matS* complexes in isolation (Dupaigne et al., 2012) (**Figure S1C**). The C-terminal tetramerization tail of MatP, however, is not visible and likely disordered, consistent with the finding that it is not required for MukBEF-related functions (Nolivos et al., 2016).

Interestingly, one of the MatP monomers forms a contact with one of the MukE monomers (**Figures 2A** and **2E**). The DNAs in ring and clamp are thus physically linked via MukE and MatP. The bridge is formed by residues between H38 and D42 in MatP and an N-terminal tail of MukE (**Figure 2E**). The latter involves residues between S2 and Q8, which are disordered in the second MukE subunit and also in previous crystal structures (Gloyd et al., 2011; Woo et al., 2009). The bridge interface is small and likely prone to dissociation, consistent with the finding that recombinantly overexpressed MukEF does not co-immunoprecipitate with purified MatP (Nolivos et al., 2016). This suggests that the bridge may have a transient role during unloading and dissociates once the reaction is complete, permitting the release of MukBEF from *matS* sites.

### The MukB joint is an interaction hub for MatP, AcpP and the hinge-proximal MukB arm

The joint of MukB is located at a central region of the complex. It is formed by an 84 amino acid insertion into the C-terminal coiled-coil strand and forms a slightly larger domain than the joints found in other SMC proteins (**Figure S3C**). The joint binds and positions MatP between the MukB arms (**Figures 2A** and **2F**). This interface is much larger than the MukE– MatP bridge, and likely provides the major binding energy for association with MatP. The joint also provides a docking site for the hinge-proximal arm, with L605 of *ν*-MukB inserted between E1137 and Y1140 of *κ*-MukB (**Figure 2G**). This likely contributes to stabilization of the elbow-folded conformation of MukBEF.

AcpP binds MukB close to the joint between R281 and F296 on the N-terminal coiled-coil helix and Y1103 and R1122 on the C-terminal helix (**Figure 2G**). Weak density protrudes from S36 of AcpP, which we have modelled as phosphopantetheine (PNS), the prosthetic group of AcpP that is covalently bound to S36 and can flip out its core upon association with binding partners (Cronan, 2014). At the *κ*-MukB binding site, PNS projects towards the space between the head-proximal arm of *κ*-MukB and the hinge-proximal arm of *ν*-MukB. The phosphate group of PNS is in contact with R839 of *ν*-MukB. PNS is modified with acyl moieties during fatty acid synthesis, and although the functional role of AcpP within the MukBEF complex is unclear, it may have a regulatory role coupling metabolism to chromosome organization (Gully et al., 2003). Due to its position near the joint–arm contact, it is possible that AcpP, perhaps via its modification state, could have an influence on the elbow-folded state of MukBEF.

The joint is situated near the heads and is thus expected to be a central conduit for conformational changes imposed by the ATPase cycle. Consistent with this idea, AcpP binding at the joint strongly increases MukBEF ATPase activity (Prince et al., 2021). Release of MukBEF from *matS* will likely require detachment from MatP, hence MatP binding at the joint seems ideal for regulation by the ATPase cycle, as will be explored below.

### Architecture of apo-MukBEF and the MukBEF dimer

In the same sample that produced reconstructions of MukBEF bound to MatP–*matS* we also observed particles with disengaged heads and neither DNA nor MatP bound (**Figure 3A**). Although the map was resolved to only 6.8 Å, which prevented determination of the nucleotide state, it was very similar to exploratory reconstructions of nucleotide-free MukBEF (**Figure S2A**). Hence, we refer to it as the ‘apo state’. The apo complex is comparable in size and shape to apo yeast condensin, with arms fully juxtaposed (**Figure S3E**). The apo clamp is more flexible because it is not held in place by ATP and DNA, but clear density was observed at lower contour levels that allowed unambiguous positioning of MukEF. The map also revealed density for the second monomer within the context of a MukBEF dimer. Further classification produced a low-resolution map for the apo dimer, from which we obtained a model by rigid body fitting (**Figure 3B**).

**Figure 3.**
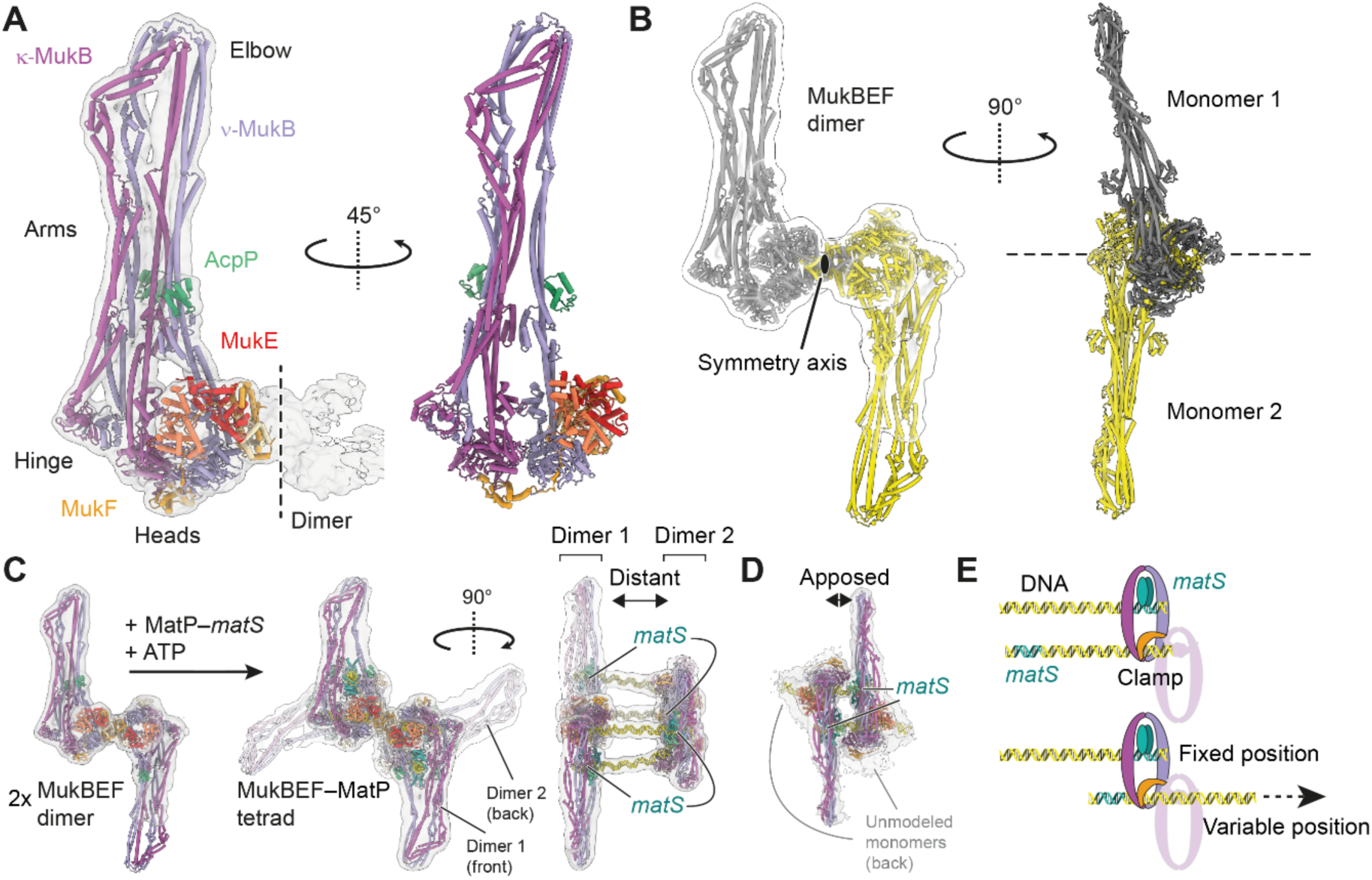
Architecture of apo-MukBEF and the MukBEF dimer. (**A**) Model of the apo-MukBEF monomer and 6.8 Å cryo-EM density at low contour level. (**B**) Model for the apo-MukBEF dimer and 13 Å cryo-EM density. (**C**) 11 Å cryo-EM density and model for two MukBEF dimers bridged by four MatP–DNA complexes (‘MukBEF tetrad’). The apo-MukBEF dimer is shown on the left. (**D**) 11 Å cryo-EM density and model for a MukBEF tetrad with closely apposed dimers. Only one monomer for each MukBEF dimer was modelled due to weak density for their partner monomers. (**E**) Schematic for variable positioning of the clamp DNA binding site, as shown in C and D. Only a single MukBEF dimer and only two of the four DNAs are shown for clarity. Related to Figures S2 and S3.

The MukBEF dimer is held together by an extensive interface between MukF’s MD and nWHD, which was observed previously by crystallography (Fennell-Fezzie et al., 2005; Woo et al., 2009). The two MukBEF monomers associate head-to-head with their MukEF subunits on the same face of the dimer. The complex thus has a front and back along its two-fold symmetry axis (**Figure 3B**).

In addition to dimers in the apo state, dimers associated with MatP and DNA were also readily resolved (**Figure 3C**). Because we positioned the *matS* site close to one end of the 80 bp DNA used for sample preparation, two dimers were able to associate with four DNAs in a ‘tetrad’ arrangement. Different classes of tetrads allowed us to assess the distance between the two dimers. We observed dimers distantly bridged by the DNA molecules (**Figure 3C**) and closely apposed (**Figure 3D**). This is explained by sequence independent binding of the clamp, which can associate with any position along the DNA double strand (**Figure 3E**). Assuming that the clamp binds DNA not only during unloading at *matS* but also during a tentative translocation reaction, it may thus step or slide along the DNA track. Since the DNAs are roughly aligned with the dimer symmetry axis, it is conceivable that DNA translocation may operate along this axis.

### Conformational changes associated with unloading

The MukB^EQ^EF–MatP–*matS* complex is blocked in ATP hydrolysis and shows a state prior to unloading. The apo form, however, lacks both MatP and DNA and therefore represents the result of a completed unloading reaction. Comparison of the two states should yield insights into conformational changes that take place during MatP-dependent exit of DNA.

Sub-classification of the cryo-EM dataset revealed additional forms of MukBEF–MatP with arms in different states of openness (**Figure 4A**). This suggests that the arms can gradually ‘zip up’, similar to what has been proposed for Smc–ScpAB based on distance measurements by electron paramagnetic resonance (Nunez et al., 2021). The most open class has arms unzipped up to the elbow, and the most closed one is the apo state. In the apo state, the heads disengage and tilt, and the joints and larynx become closely juxtaposed (**Figure 4B**). This occludes the binding sites for both MatP–*matS* and the clamped DNA and strongly supports the idea that ATP hydrolysis promotes dissociation from MatP–*matS* and DNA unloading.

**Figure 4.**
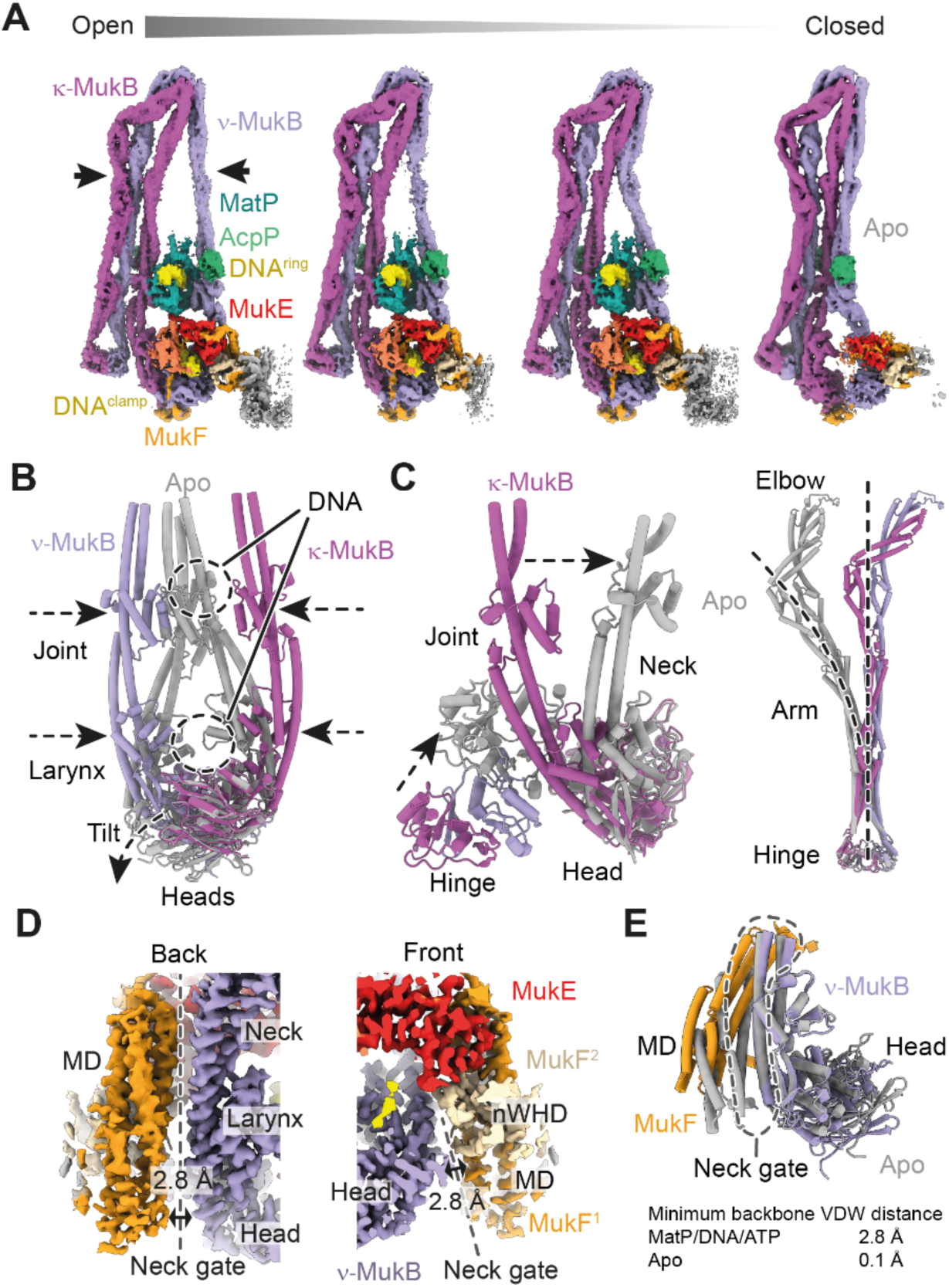
Conformational changes associated with release of MatP/DNA/ATP. (**A**) Cryo-EM densities for the MukBEF–MatP–DNA complexes with different arm conformations. Density of apo-MukBEF with fully closed arms is shown on the right. (**B**) Blocking of MatP and DNA binding sites at the MukB joint and larynx. Structures were superimposed on the ATPase domains. MatP/DNA/ATP-bound conformation is shown in color, apo conformation in grey. (**C**) Conformational change at the MukB neck/hinge interface (left) and at the hinge-proximal arm (right). Structures were superimposed on the ATPase domain (left) or the hinge (right). (**D**) Cryo-EM density at the neck gate in the MatP/DNA/ATP bound state. The solvent accessible cleft between MukB and MukF is indicated by a double arrow. (**E**) Superimposition of the neck gate in apo and MatP/DNA/ATP bound states (top). Minimum backbone VDW distances of the interface are given (bottom). See also Figures S2 and S3.

Conformational changes associated with the release of nucleotide and DNA propagate through the whole complex and can be observed even in the hinge-proximal coiled coil (**Figure 4C**). The hinge follows the tilting *κ*-MukB neck during DNA unloading, and the hinge-proximal arm changes from a straightened conformation to a strongly curved one. This indicates that the arm stores parts of the binding energy provided by ATP, MatP and DNA as elastic energy, similar to a spring. This energy may be harnessed to expel MatP and DNA after ATP hydrolysis.

In cohesin, DNA unloading proceeds via opening of the interface between the kleisin Scc1 and the *ν*-SMC Smc3 (Chan et al., 2012; Muir et al., 2016; Murayama and Uhlmann, 2015). In MukBEF, the corresponding interface, which we name ‘neck gate’, is formed by the neck of *ν*-MukB, and the MD of one MukF together with the nWHD of the second MukF (**Figures 4D**). Intriguingly, the neck gate is opened by a narrow cleft along the interface in the MatP/DNA bound structure but is closed in the apo structure (**Figures 4D** and **4E**). The minimum backbone Van-der-Waals (VDW) distance is 2.8 Å across the interface in the MatP/DNA bound state, which is reduced to 0.1 Å minimum backbone VDW distance in the apo structure. Although the cleft is too narrow for DNA to pass through, it may represent a step towards full opening of the neck gate.

### MukBEF adopts a folded conformation *in vivo*

MukBEF adopts an elbow-folded conformation, at least in its apo state and when bound to ATP, MatP and DNA. However, the elbow has also been crystallized in an extended conformation, suggesting that MukBEF may convert to extended rods or fully open rings with disengaged arms (Bürmann et al., 2019) (**Figure 5A**). The hinge-proximal arms are in a closed rod conformation in our structures, similar to those of other SMC complexes (**Figure S3F**), but a conformation compatible with open rings has been observed by crystallography (Li et al., 2010) (**Figure S3G**). Additional support for the existence of open rings comes from rotary shadowing electron microscopy experiments (Matoba et al., 2005). We thus decided to clarify whether the elbow-folded conformation is abundant *in vivo*, and whether it may be controlled by the ATPase cycle of MukBEF. To accomplish this, we probed the conformation of endogenous *E. coli* MukBEF by site-specific cysteine cross-linking.

**Figure 5.**
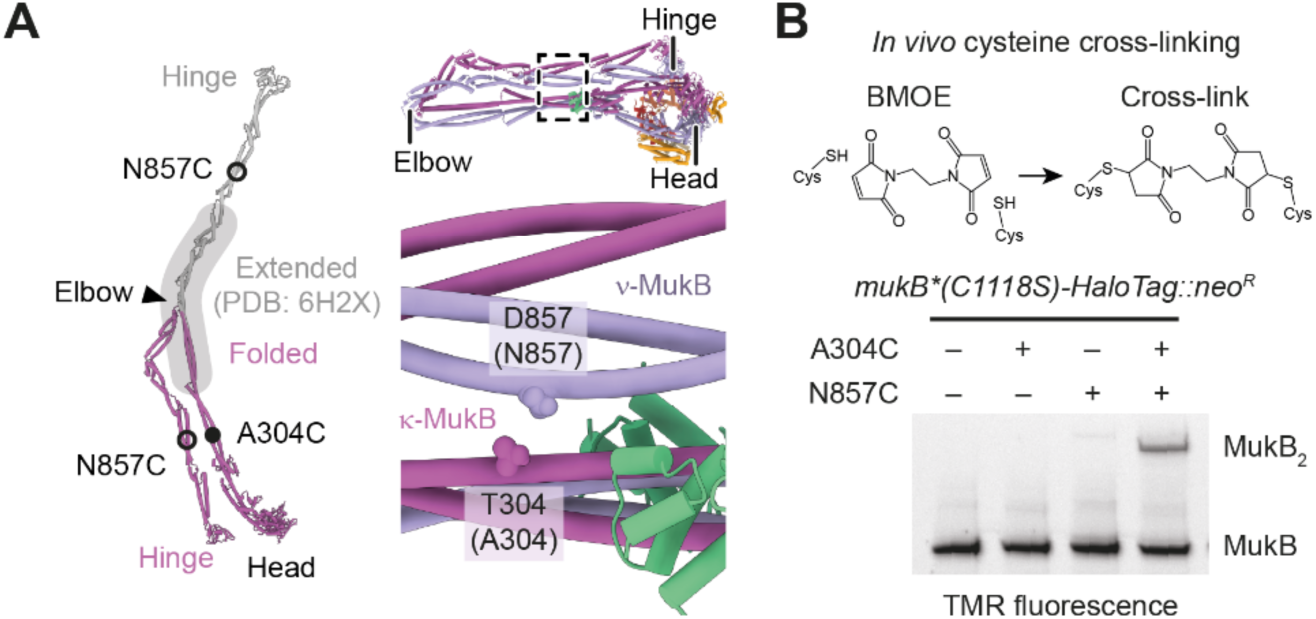
Detection of arm folding *in vivo*. (**A**) Residues employed as sensors for the folded conformation. The folded conformation and a tentative extended conformation based on the structure of the extended elbow (PDB: 6H2X) are shown on the left. A close-up on the *P. thracensis* structure is shown on the right. Corresponding *E. coli* residues are in parentheses. (**B**) BMOE reaction scheme (top) and BMOE mediated *in vivo* cysteine cross-linking of *E. coli* strains carrying sensor cysteine mutations (bottom). Reaction products were detected by SDS-PAGE and in-gel fluorescence using a TMR fluorophore bound to MukB-HaloTag. See also Figures S4 and S5.

To introduce multiple point mutations spread across the 8 kb chromosomal *mukFEB* locus, we used a derivative of REXER (replicon excision for enhanced genome engineering through programmed recombination) (Wang et al., 2016). In addition to introducing mutations A304C and D857C into MukB to probe the folded conformation (**Figure 5A**) we introduced C1118S, which ablated weak background cross-linking with a small protein, possibly AcpP (**Figure S4**). Next, we treated the *E. coli* cells with bismaleimidoethane (BMOE), which rapidly *in vivo* cross-links closely spaced thiols such as cysteine sidechains. We then detected reaction products labelled with HaloTag-tetramethylrhodamine (TMR) by SDS-PAGE and in-gel fluorescence (**Figure 5B**). Residues A304C and D857C cross-linked specifically with close to 50 % efficiency, demonstrating that the folded conformation exists *in vivo*. The reaction efficiency was comparable to that of constitutive interfaces (see below), indicating that the folded state is abundant.

Next, we generated ATPase mutant strains S1366R (MukB^SR^, blocking head engagement), D1406A (MukB^DA^, blocking ATP binding) or E1407Q (MukB^EQ^, blocking ATP hydrolysis) (Woo et al., 2009) (**Figure S5A**). As expected, all mutations conferred a *mukB* null phenotype, characterized by an inability to grow on rich media at 37 °C. To probe the effect of the mutations on the ATPase cycle, we then introduced G67C, which is located at the top of the MukB head and close to its symmetry mate in the second MukB (**Figure S5A**). This residue changes distance upon head engagement and should respond to alterations in the ATPase cycle. The residue cross-linked with similar efficiencies in WT and the ATP binding mutant MukB^DA^, with 22 ± 1 % and 21 ± 2 %, respectively. The cross-linked fraction was increased to 30 ± 1 % in MukB^EQ^, indicating enhanced head engagement (**Figure S5B**). This confirms that the assay is able to detect conformational changes *in vivo*, and also suggests that the heads of wild-type MukBEF are mostly disengaged.

We then probed for elbow folding in MukB^SR^, MukB^DA^ and MukB^EQ^ strains which resulted in similar reaction efficiencies to WT (**Figure S5C**). These findings suggest that the elbow folded state of MukBEF is not controlled by steps preceding ATP hydrolysis, consistent with our structural data.

### Clamp and ring compartment entrap separate segments of a DNA loop *in vivo*

MukBEF–MatP entraps DNA within its ring and clamp compartments. This implies that, in the context of a circular chromosome, DNA would have to enter through one or more entry gates. For biochemical preparation of the cryo-EM sample, however, any loading or partial unloading reactions were bypassed using linear DNA and an ATPase-deficient mutant. As an additional caveat, linear DNA prevents determination of the DNA connectivity that would occur in a physiological context. For example, DNA in the ring and DNA in the clamp may originate from the same or from different chromosomes. Hence, we decided to map the DNA binding topology of MukBEF *in vivo*.

We adapted an assay that measures chromosome entrapment by chemically circularized protein complexes for the use in *E. coli* (Vazquez Nunez et al., 2019; Wilhelm et al., 2015). In this assay, covalent circularization of a protein compartment around chromosomal DNA preserves DNA association after denaturation of DNA-binding surfaces (**Figure 6A**). Guided by structural information, we designed cysteine cross-links at the cap (S143C in MukB and Q412C in MukF), the neck (K1246C in MukB and D227C in MukF), and the hinge (C730 and R771C in MukB) to probe DNA entrapment in the ring compartment (**Figure 6B** and **S6A**). We also combined cap and neck cysteines with the G67C head cysteine to probe entrapment in the clamp (**Figures 6B** and **S5A**). In addition, head and hinge cysteines were combined to probe entrapment in the ‘frame’ compartment, which is the union of ring and clamp.

**Figure 6.**
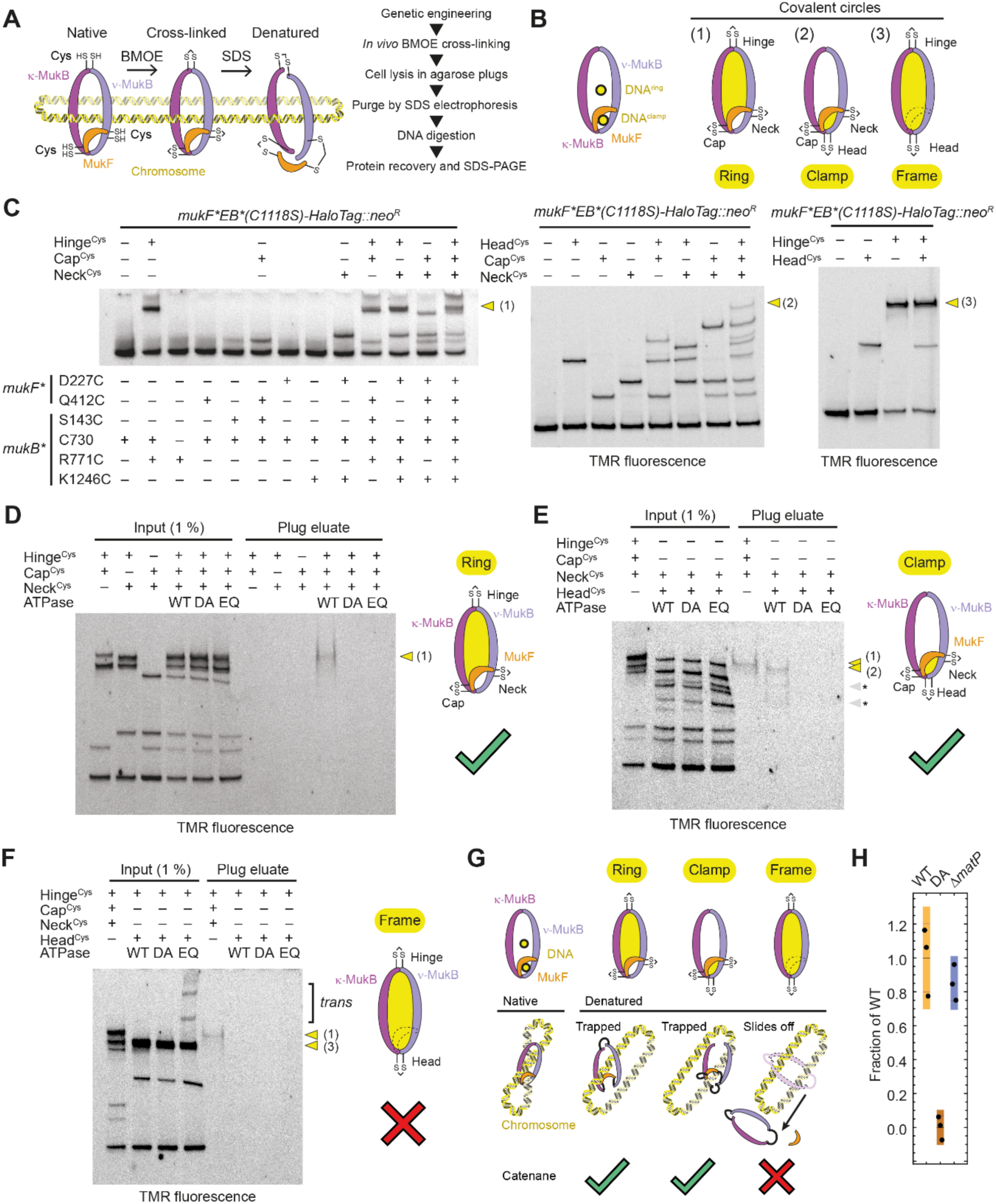
Mapping of DNA binding topology *in vivo*. (**A**) Principles (left) and workflow (right) of the chromosome entrapment assay in agarose plugs. (**B**) Combinations of cross-links used for probing DNA entrapment in ring, clamp and frame compartments. Hinge cross-link: C730 and R771C in MukB; cap cross-link: Q412C in MukF and S143C in MukB; neck cross-link: D227C in MukF and K1243C in MukB; head cross-link: G67C in MukB (Figures S5A and S6A). (**C**) Combinatorial cross-linking for identification of reaction species. Combinations: Hinge, cap, and neck cross-links (left); cap, neck, and head cross-links (middle); head and hinge (right). C730 was mutated to serine when indicated by ‘– ‘. Cells were grown to stationary phase. Detection as in Figure 5B. (**D**) Chromosome entrapment in MukBEF with covalently closed ring compartment. Input and agarose plug eluate are shown. Detection as C. The circular species is retained only in WT ATPase cells. DA, D1406A (blocks ATP binding); EQ, E1407Q (blocks ATP hydrolysis). ATPase is WT if not indicated otherwise. (**E**) Chromosome entrapment in MukBEF with covalently closed clamp compartment. As in D. Species that have undergone chemical cross-link reversal during sample preparation are marked with asterisks. (**F**) Chromosome entrapment in MukB with covalently closed frame compartment. As in D. Species produced by higher oligomers in EQ mutants (*trans* cross-links) are indicated. (**G**) Structure-based topological interpretation of the entrapment reactions. Only the frame species can slide off DNA, because it does not form a protein/DNA catenane. (**H**) Chromosome entrapment in the ring in the absence of MatP. Entrapment signal relative to WT is shown for biological triplicates. Black lines indicate means, purple lines indicate standard deviations, and colored bars indicate 95 % credible intervals. See also Figures S5 and S6.

Combinations of cysteines were introduced into the endogenous *mukFEB* locus, cells were treated with BMOE, and reaction species were identified by SDS-PAGE and in-gel fluorescence (**Figure 6C**). Cross-linking was specific and efficient at all sites. For some reaction products, species were not completely resolved from each other due to high molecular weight, identical mass and mere shape differences. However, depletion of precursors in expected ratios indicated successful multi-site reactions in all cases. For example, the MukB species cross-linked at the heads was reduced from 29 % to 12 % when combined with the hinge cross-link. This corresponds to a reduction by 60 % and is in excellent agreement with the 62 % cross-linking efficiency observed for the hinge.

First, we tested entrapment in the ring compartment using cap, neck, and hinge cysteine pairs. We treated cells with BMOE, lysed them in agarose plugs to protect chromosomal DNA from shearing and subjected plugs to electrophoresis in the presence of 0.1% sodium dodecyl sulphate (SDS). This denatures and extracts proteins and only retains cross-linked species that have been circularized around DNA. We then digested chromosomal DNA and isolated the eluted proteins. A cross-linked high-molecular weight MukB species was retained only when cap, neck and hinge interfaces all contained cysteine pairs, indicating that it is the covalently circularized MukB_2_F (**Figure 6D**). The species was not retained from MukB^DA^ or MukB^EQ^ strains, which suggests that DNA entrapment critically depends on ATP hydrolysis.

Next, we tested for DNA entrapment in the clamp compartment using cap, neck, and head cysteines (**Figure 6E**). The cross-linked MukB species corresponding to the circularized clamp was isolated as the major band, accompanied by small amounts of protein presumably resulting from chemical cross-link reversal during the protein isolation procedure (Shen et al., 2012; Wilhelm and Gruber, 2017). The retained amount was similar to that of the ring compartment species, consistent with the notion that both clamp and ring entrap DNA simultaneously. Protein was not retained from MukB^DA^ or MukB^EQ^ strains. This finding indicates that DNA entrapment in the clamp, as in the ring, depends on ATP hydrolysis *in vivo*.

Next, we tested for DNA entrapment in the frame compartment using the combination of head and hinge cysteines (**Figure 6F**). Head and hinge cross-linked MukB was neither detected in eluates from WT, MukB^DA^ or MukB^EQ^ strains. This and the above two findings are best explained by the notion that ring and clamp each entrap different strands of the same loop. Cross-linking the frame around this loop allows DNA to slip out of the covalent protein circle, because no protein–DNA catenane is formed (**Figure 6G**).

Taken together, these results fully support entrapment of a loop as suggested by the structure (**Figures 1H, 6G** and **S6B-S6C**). Because no DNA entrapment is detected in the frame compartment, and signals for clamp and ring are similar, most if not all complexes with DNA in the clamp must have DNA catenated with the ring, and vice versa. The results are incompatible with entrapment in only ring or clamp, with entrapment of sister chromosomes, or with a loop axis running parallel to the plane of the ring (**Figure S6C**). Any of these forms would lead to catenation with the frame compartment.

Finally, we investigated whether chromosome entrapment was dependent on MatP. Deletion of the *matP* gene had little if any effect on DNA inside the MukBEF ring (**Figure 6H**). We suggest that DNA entrapment is a more general feature of MukBEF and does not exclusively occur during unloading at *matS* sites.

## Discussion

### The double-locked loop

The structure of MukBEF bound to MatP–*matS* revealed the simultaneous entrapment of two DNA double helices, topologically separated into ring and clamp compartments. We name this configuration the ‘double lock’ (**Figure 7A**). The DNA crossing angle in the MatP-bound double lock indicates that a DNA loop passes through MukBEF with the loop long axis perpendicular to the ring plane. Topological mapping *in vivo* supports this notion.

**Figure 7.**
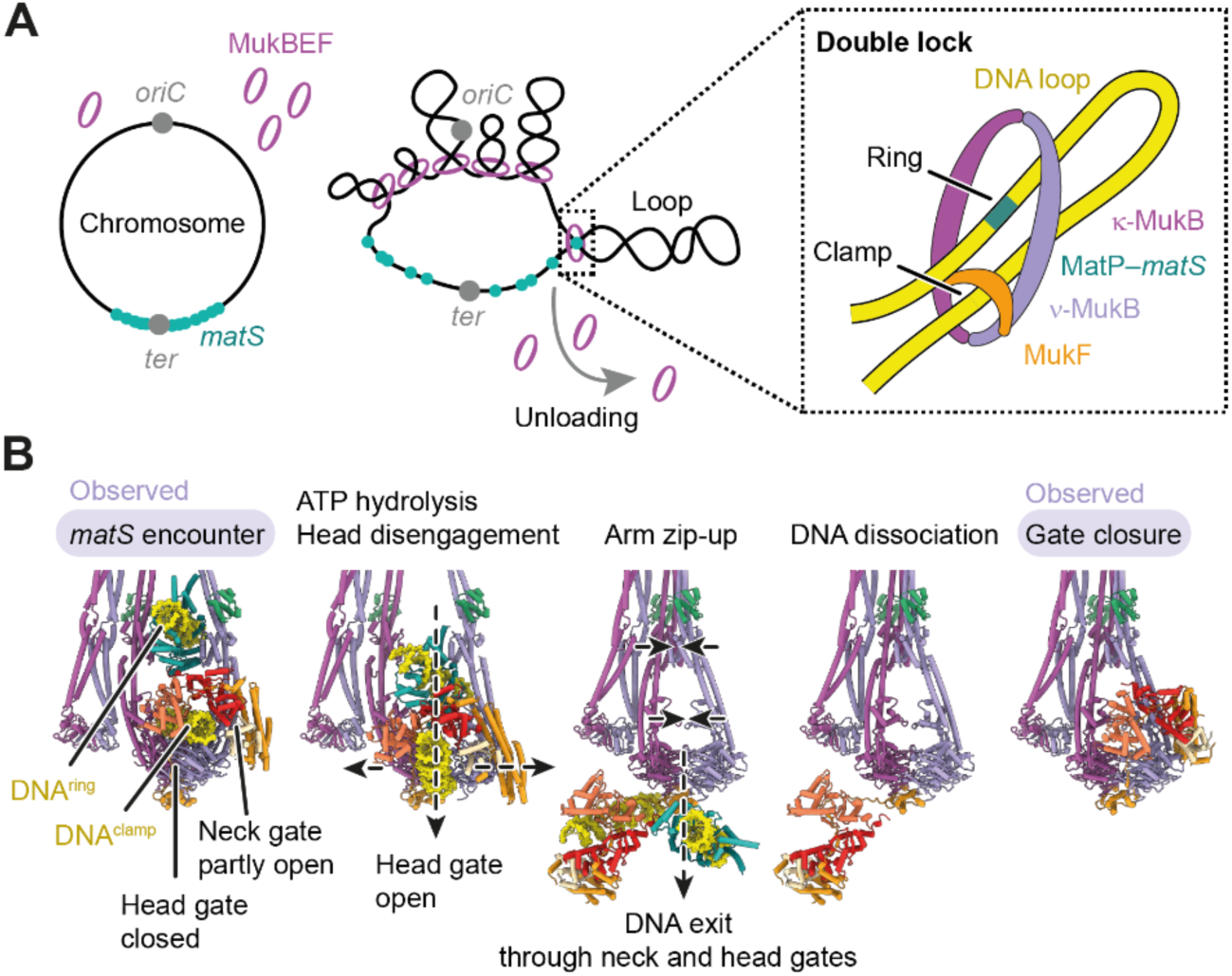
Model for DNA binding and unloading at *matS* sites. (**A**) Schematic for association of MukBEF with MatP–*matS* in the *Ter* macrodomain. MukBEF organizes the chromosome into loops. Upon invasion of *Ter*, MukBEF encounters MatP–*matS* and unloads via the double-lock topology in the context of a plectonemic loop. (**B**) Model for unloading of DNA. A MatP–*matS* encounter is followed by ATP hydrolysis and opening of the head gate to permit exit of the clamped DNA. *matS* DNA follows through neck and head gates, facilitated by the bridge between MatP and MukE. Arm zip-up prevents reversal, and the neck gate closes after DNA/MatP dissociation. States observed experimentally are highlighted in purple.

DNA entrapment by MukBEF occurs mainly outside of the MatP–*matS* context. Because localization of MukBEF to chromosomal foci is largely independent of MatP, and focal MukBEF turns over in an ATP hydrolysis dependent manner (Badrinarayanan et al., 2012; Mäkelä and Sherratt, 2020b; Nolivos et al., 2016), it is conceivable that unloading via the double lock happens throughout the chromosome. At *matS* sites, MatP may enhance the positioning of DNA within the ring for an efficient unloading reaction.

How abundant is the double lock? The fraction of cross-linked species retained by the entrapment assay is low, raising the possibility that the double lock is a sparsely populated state. However, the relative signal of the assay will depend on factors such as preservation of chromosomal DNA and the fraction of MukBEF complexes that is loaded. The assay therefore likely underestimates abundance by an unknown and possibly large factor, and may thus not be suited for its quantification. In other words, the data support the existence of the double lock, but do not necessarily reveal its incidence. What the experiments do suggest, however, is that a large fraction of MukBEF with a loaded clamp is in the double lock configuration. Hence, transactions that involve DNA binding inside the clamp may largely progress via the double lock. The same statement applies to DNA entrapment inside the ring. DNA transactions that involve catenation with the ring may predominantly progress via the double lock.

### The role of MukBEF dimerization

The architecture of the MukBEF dimer permits binding of two double-locked loops, whereby the loop axes are parallel, and monomers are arranged side-by-side. If loop extrusion by MukBEF monomers was an asymmetric process, similar to what has been observed for monomeric condensin (Ganji et al., 2018), such side-by-side coupling could symmetrize the overall process. Highly asymmetric loop extrusion is considered a hindrance for physiological chromosome folding and may be resolved using a dimerization mechanism (Banigan and Mirny, 2019). We suggest that the arrangement of the MukBEF dimer is well suited to implement symmetric loop extrusion.

### DNA entrapment in MukBEF and other SMC complexes

MukBEF does not entrap sister DNAs or loops with a long axis perpendicular to the one proposed (**Figure S6C**). However, the low sensitivity of our assay may have precluded detection of rare species. Our experiments also do not exclude formation of ‘pseudo-topological’ loops or ‘non-topological’ loops, which may form in addition to the double lock. The former are established by threading DNA through the same compartment twice, constituting a protein–DNA rotaxane instead of a catenane, whereas the latter do not thread through the complex at all but bind to its outer surface (**Figure S6C**). These structures may be involved in loop extrusion by cohesin (Davidson et al., 2019; Srinivasan et al., 2018), but their binding by MukBEF is hypothetical. If ‘pseudo-topological’ loops exist, however, they will need to be accommodated in the ring and not in the clamp due to space limitations (**Figure S6B**). Whereas the clamp is highly constrained by a short and compact MukEF and can thus accommodate only a single DNA, the ring would be able to embrace multiple DNAs and could even enlarge its capacity by extending the elbow and fully opening the arms.

Cysteine cross-linking has been used to map DNA entrapment in other SMC complexes (Chapard et al., 2019; Gligoris et al., 2014; Haering et al., 2008; Vazquez Nunez et al., 2019; Wilhelm et al., 2015). Cross-linking at cap, neck and hinge of cohesin and Smc–ScpAB (also designated as ‘SK’ cross-links) retains either complex on DNA after denaturation. When cross-linked at cap, neck and juxtaposed heads (also designated as ‘JK’ cross-links), DNA association is also maintained. Entrapment is not observed when cysteines in hinge and juxtaposed heads are combined (also designated as ‘JS’ cross-links). These patterns are topologically equivalent to the ones determined here. Hence, the MukBEF structure may provide an attractive interpretation for these observations, namely that cohesin and Smc–ScpAB may be able to entrap DNA in a manner similar to the double-lock configuration.

Interestingly, the head cross-links used for cohesin and Smc–ScpAB can capture a ‘head juxtaposed’ state, which is different from engaged heads in the corresponding ATP hydrolysis mutants. MukBEF heads are mostly disengaged *in vivo* as judged by our cross-linking experiments, and the G67C cross-link at the heads used for topology mapping can capture the disengaged state. It is therefore possible that DNA may be retained in the clamp even after ATP hydrolysis.

### Implications for chromosomal turnover of MukBEF

Both ring and clamp of MukBEF fully encircle DNA. This raises the question of how DNA enters and exits these compartments. The structure of MukBEF–MatP–*matS* shows a state before DNA release, whereas the apo structure shows the state after release. The latter also represents the state before DNA has entered the complex. Models for chromosomal turnover of MukBEF will have to fit these observations.

We envision that unloading of *matS* from the ring is coupled to unloading of DNA from the clamp (**Figure 7B**). For the clamped DNA, the interface between the heads is a prime candidate for an exit gate because its formation is regulated by ATP binding and hydrolysis. Upon ATP hydrolysis and phosphate/nucleotide release, heads disengage and dissociate their DNA binding surfaces. Clamped DNA may then be able to exit via the cleft between disengaging heads. For this to happen, the MukF linker, which also seals the head gate, needs to detach from the *ν*-MukB head. MukEF can stay bound to and move along with the DNA because the neck gate is open, which allows MukEF to reposition in relation to the *ν*-MukB head. Concurrently, a deformation at the joints releases MatP, which stays associated with MukEF via the MatP–MukE bridge. Next, the head proximal arms of MukB zip up and occlude the binding sites for MatP and DNA at the joints and heads, respectively. Release of energy stored as deformations in the hinge-proximal arm will reinforce this process. MatP and DNA are ejected and are free to dissociate from the complex. Finally, closure of the neck gate reverts MukBEF to its apo form. We note that the proposed mechanism would not strictly depend on MatP–*matS* but could also eject ‘free’ DNA from the ring compartment. Its driving force comes from relaxation of MukB into the apo conformation, whereas MatP is a structural element that ensures ideal positioning of DNA close to the exit gate.

The proposed unloading model, which is purely based on structural data, is attractive for several reasons. First, it explains how the process is regulated by ATP hydrolysis, namely by opening the head gate and blocking the binding sites for MatP and DNA. Second, it explains how the process is enhanced at *matS* sites. Third, the model designates the neck gate as the topological exit gate of the tripartite ring. The equivalent interface in the distantly related cohesin is the exit gate of this complex (Chan et al., 2012; Muir et al., 2020; Murayama and Uhlmann, 2015). We suggest that unloading via the neck gate is a widely conserved activity of SMC complexes.

An interaction of MatP with the MukB hinge, at least in the absence of DNA, has been reported (Fisher et al., 2021; Nolivos et al., 2016) but is not seen in our structures. It is conceivable that this occurs at a different stage during unloading and could suggest subunit and DNA transport within the complex. Interestingly, MukB^EQ^ can associate with *matS* in cells (Nolivos et al., 2016), but DNA entrapment is not detected in our assay. This may point towards a state which binds MatP but does not entrap DNA.

In the light of our structures, DNA entry into MukBEF appears enigmatic, as is the case for other SMC complexes. The joint, which has emerged as a central region regulated by the ATPase, is required for recruitment of Smc–ScpAB to its loading factor ParB (Gruber and Errington, 2009; Minnen et al., 2016). ParB is typically not present bacteria that use MukBEF, and no substitute loading factor has been identified. Importantly, loading of MukBEF does not depend on MatP, and a reverse and MatP-independent version of the unloading mechanism proposed above would have to work against a large entropic barrier. This rather unlikely pathway would also depend on ATP binding only, whereas loading *in vivo* requires nucleotide hydrolysis. We suspect that chromosomal loading and unloading of MukBEF are achieved by considerably different means.

### Outlook

Here we have determined the structure of MukBEF both in its apo state and in a MatP/DNA/ATP bound form, providing molecular insight into how MukBEF is released from chromosomes. This led to the finding that MukBEF can entrap DNA loops in a double-lock configuration, which links two important topological concepts, namely DNA entrapment inside the ring and inside the clamp compartment. Our work opens up the questions of how the double-locked loop is established, and how MukBEF operates on this and possibly other types of DNA structures. We anticipate that biochemical reconstitution of the process by which MukBEF organizes chromosomes, coupled to structural analysis, will further our understanding of chromosome folding in bacteria and beyond.

## Data availability

EM density maps have been deposited in the EMDB (EMD-12656, EMD-12657, EMD-12658, EMD-12659, EMD-12660, EMD-12662, EMD-12663, EMD-12664). Coordinates have been deposited in the PDB (7NYW, 7NYX, 7NYY, 7NYZ, 7NZ0, 7NZ2, 7NZ3, 7NZ4). See also **Table S1**. All other data will be available upon request.

## Material & Methods

### Protein production and purification

Wild-type *P. thracensis* MukBEF (NCBI accession identifiers WP_046975681.1, WP_046975682.1, and WP_046975683.1) was produced from a polycistronic expression construct assembled into a pET28 based backbone by Golden Gate cloning (Engler et al., 2008) (pFB403). The construct contained a His_6_-SMT3 tag fused to residue 1 of MukB which allowed affinity purification and scar-less tag removal by hSENP2 protease (Butt et al., 2005). The complex was produced in BL21-Gold(DE3) by autoinduction (Studier, 2005) in ZYP-5052 media at 24 °C. All purification steps were carried out at 4 °C. 15 g of cells were resuspended in 90 mL of IMAC buffer (50 mM Tris, 300 mM NaCl, 40 mM imidazole, 1 mM TCEP, pH 7.4 at room temperature (RT)) supplemented with protease inhibitor cocktail (Roche) and Benzonase (Merck) and lysed at 172 MPa in a high-pressure homogenizer. The lysate was cleared by centrifugation at 96,000 x g for 30 min, passed through a 0.45 μm filter, and incubated for 30 min with 25 mL Ni-NTA agarose (Qiagen) equilibrated in IMAC buffer. The resin was packed into a gravity flow column and washed with 3 x 50 mL IMAC buffer, then resuspended in 25 mL IMAC buffer containing 1 mg GST-hSENP2 and incubated for 1 h on a roller. Eluate was collected and pooled with a 12.5 mL wash using IMAC buffer, diluted with 18.8 mL buffer Q (10 mM Tris, pH 7.4 at RT), passed through a 0.22 μm filter and applied to a 20 mL HiTrap Heparin HP column (GE Healthcare). MukBEF was largely found in the flowthrough and was applied to a 5 mL HiTrap Q HP column (GE Healthcare). The column was washed with 2 column volumes (CV) of 10 mM Tris, 200 mM NaCl, 1 mM TCEP, pH 7.4 at RT, and protein was eluted with a 20 CV linear gradient from 200 mM NaCl to 1 M NaCl in buffer Q. MukBEF eluted at about 450 mM NaCl, was concentrated to 0.5 mL on a Vivaspin 20 MWCO 30 filter (Sartorius) and was injected into a Superose 6 Increase 10/300 column (GE Healthcare) in buffer H200 (20 mM Hepes, 200 mM NaCl, 1 mM TCEP, pH 7.3 at RT). Peak fractions were pooled, concentrated to 8.2 mg/mL on a Vivaspin 2 MWCO 30 filter, aliquoted, frozen in liquid nitrogen and stored at -80 °C until use. Protein stoichiometry was estimated by SDS-PAGE and Coomassie staining as MukB_2_E_4_F_2_–AcpP_2._

We were unable to establish polycistronic MukB^EQ^EF expression constructs, likely due to toxicity in the cloning host. Therefore, we cloned His_6_-SMT3-MukB^EQ^ and MukEF on two separate expression constructs, pFB485 and pFB486, whereby the His_6_-SMT3-MukB^EQ^ construct was always propagated at 22 °C. Proteins were separately produced as above, with His_6_-SMT3-MukB^EQ^ production at 22 °C. Cell pellets of both strains (15 g each) were mixed in 180 mL IMAC buffer, and the complex was purified as described above, except that 1 mM EDTA (pH 7.4 at RT) was added before application to the Heparin column with the intention to improve dissociation of potentially co-purifying nucleotides. Estimated stoichiometry was MukB^EQ^ E_4_F_2_–AcpP_2._

MukB^EQ^ was purified as above except for omission of the Heparin step. Protein was concentrated to 8 mg/mL. Estimated stoichiometry was MukB^EQ^ –AcpP_2._

*P. thracensis* MatP (NCBI accession identifier WP_046976581.1) was cloned without tag (pFB469) and expressed as above. The untagged protein bound tightly to IMAC resin. Extract was prepared as above and was passed through a 5 mL HisTrap HP column (GE Healthcare). The column was washed with 10 CV of IMAC buffer, and the protein was eluted with IMAC buffer containing 270 mM imidazole. The eluate was diluted with an equal volume of buffer Q, passed through a 0.45 μm filter and applied to a 5 mL HiTrap Q HP column. MatP was largely found in the flowthrough, which was then loaded on a 5 mL HiTrap SP HP column (GE Healthcare). The column was washed with 2 CV buffer Q containing 150 mM NaCl and eluted with a 20 CV linear gradient from 150 mM to 1 M NaCl in buffer Q. The protein eluted at around 400 mM NaCl. Peak fractions were pooled, concentrated in a Vivaspin 20 MWCO 10 filter (Sartorius) to 4.8 mg/mL, aliquoted, frozen in liquid nitrogen and stored at -80 °C.

All protein concentrations were determined by absorbance at 280 nm using theoretical absorption coefficients. Annotated sequences of expression constructs are provided in **Data S1**.

### *matS* sites and electrophoretic mobility shift assay (EMSA)

Initial binding experiments indicated that the affinity of *P. thracensis* MatP for consensus *E. coli matS* <u>GTGAC</u>ATT<u>GTCAC</u> (palindrome underlined) was about an order of magnitude lower than reported for other MatP proteins (Dupaigne et al., 2012; Mercier et al., 2008). To identify alternative *matS* sites, we mapped all sites with edit distance up to 2 within the palindromic region onto the *P. thracensis* chromosome (NCBI accession identifier CP011104.1) using Wolfram Mathematica, and ranked them by median distance to the replication terminus predicted by cumulative GC skew (Lobry, 1996). This identified <u>GTTAC</u>NNN<u>GTAAC</u> as an abundant site with 9 occurences (**Figure S1A**) which was further characterized by EMSA.

6-Carboxyfluorescein (6-FAM) labelled DNA oligonucleotides *matS1* (annealed from single stranded oligonucleotides [6-FAM]CACT<u>GTGACATTGTCAC</u>GGCA and TGCC<u>GTGACAATGTCAC</u>AGTG; *E. coli* consensus *matS* underlined) or *matS2* ([6-FAM]CACT<u>GTTACAGTGTAAC</u>GGCA and TGCC<u>GTTACACTGTAAC</u>AGTG; *P. thracensis* candidate site underlined) at a final concentration of 2 nM in buffer H30 (20 mM Hepes, 30 mM NaCl, 1 mM TCEP, pH 7.3 at RT) were titrated with MatP and incubated for 5 min at RT. Samples were resolved on a 6 % DNA Retardation gel (Thermo Fisher Scientific) with 0.5x TBE running buffer at 100 V for 60 min at 4 °C. Gels were scanned on a Typhoon TLA9000 with Cy2 setup.

Quantification of bands was performed with Wolfram Mathematica using moving median estimation of background signal. Data were fit by a rate equation model at equilibrium in an arbitrary time domain, parametrized by dissociation rate k_d_, baseline and asymptote, and with the association rate k_a_ as an arbitrary model constant. The equilibrium dissociation constant K_d_ was determined as k_d_/k_a_ which is independent of the time domain. This approach is easily extendable to reaction models where analytical solutions or approximations are difficult to derive and can be adapted to time resolved detection methods for determination of true rate constants.

### Cryo-EM sample preparation

Optimal protein ratio for formation of MukBEF dimers was estimated by titrating 375 nM MukB_2_E_4_F_2_–AcpP_2_ with MukB_2_–AcpP_2_ in buffer H200 (20 mM Hepes, 200 mM NaCl, 1 mM TCEP, pH 7.3 at RT), followed by size exclusion chromatography on a Superose 6 Increase 3.2/300 column (GE Healthcare) mounted on an ÄKTA Ettan (GE Healthcare). An equimolar mixture shifted most protein to the dimer fraction, with some residual material in MukB_2_E_4_F_2_– AcpP_2_ and MukB_2_–AcpP_2_ fractions.

MukB^EQ^ E_4_F_2_–AcpP_2_ and MukB^EQ^ –AcpP_2_ were mixed at 10 μM each in 4.8 μL buffer H200 (20 mM Hepes, 200 mM NaCl, 1 mM TCEP, pH 7.3 at RT) and incubated for 10 min on ice. Then, 0.48 μL of a 100 μM stock of DNA in water (annealed from oligonucleotides CTCGCCTGTAAAGTAGGCATTAGTTGTTCGTAGTGCTCGTCTGGCTCTGGATTACCCGCCACT<u>GTTAC ATTGTAAC</u>GGCA and TGCC<u>GTTACAATGTAAC</u>AGTGGCGGGTAATCCAGAGCCAGACGAGCACTACGAACAACTAATGCCT ACTTTACAGGCGAG; *matS* sequence underlined) and 1.8 μL of MatP_2_ (26.8 μM stock in buffer H200) were added. After 5 min on ice, 4.9 μL of buffer H30 (20 mM Hepes, 30 mM NaCl, 1 mM TCEP, pH 7.3 at RT) were added and the sample was passed over a Zeba Micro Spin 7K MWCO column (Thermo Fisher Scientific) equilibrated in buffer H30 containing 1 mM ATP, 2 mM MgCl_2_ and 0.05 % (w/v) *β*-octyl glucoside. The sample was incubated for 10 min at RT, then for 30 min on ice.

Cryo-EM grids were prepared as follows. UltrAuFoil (Russo and Passmore, 2014) R2/2 200 mesh grids (Quantifoil) were glow discharged for 15 s at 30 mA in a Quorum SC7620, and 2.5 μL sample were applied to a freshly glow discharged grid mounted in a TFS Vitrobot Mark IV, equilibrated at 4 °C and 100 % humidity. Grids were immediately blotted at blot force -15, 2-4 s blotting time, no drain time and plunge frozen in liquid ethane. Grids were stored in liquid nitrogen until use. Sample screening and optimization was performed on TFS Tecnai F20, Polara and Glacios microscopes.

### Cryo-EM data collection

Data was collected on multiple grids over three sessions on a TFS Titan Krios with X-FEG emitter at 300 kV, equipped with a Gatan K3 detector operating in counting mode and a Gatan Quantum energy filter with 20 eV slit width centered on the zero-loss peak. Datasets 1 and 2 were collected with hardware binning at 1.07 Å calibrated pixel size. Dataset 3 was acquired without hardware binning at 0.535 Å calibrated pixel size. Movies were acquired in SerialEM (Mastronarde, 2005) at three areas per hole, using image shift with beam tilt compensation to collect 9 holes per stage movement. Target defocus was -1 to -3 μm, total electron fluence was 40 e^-^/A^2^ collected over 2.5 s and fractionated into 40 frames.

### Cryo-EM data analysis

An overview of the data analysis workflow is shown in **Figure S2**. Motion correction and dose weighting was performed in RELION (Scheres, 2012) with one patch per micrograph and on-the-fly gain correction. Super-resolution data of dataset 3 were binned by a factor of 2. The contrast transfer function (CTF) was fitted with CTFFIND4 (Rohou and Grigorieff, 2015). Particle picking was performed with crYOLO (Wagner et al., 2019). All further processing was done in RELION.

An initial model of MukBEF was reconstructed from an exploratory dataset collected on apo- MukB_2_E_4_F_2_–AcpP_2_ (**Figure S2A**). Particles were picked with a custom trained crYOLO model and subjected to 2D classification. Particles from good classes were used for *ab initio* reconstruction. The model was manually sculpted at to remove obvious artefacts, and iteratively improved by 3D classification, auto-refinement, sculpting, optimization of the crYOLO model and repicking.

MukB^EQ^EF–MatP–DNA particles were picked from dataset 1 using the apo-MukBEF crYOLO model. Using the apo-MukBEF reconstruction as a starting model, an initial model for MukB^EQ^EF–MatP–DNA was obtained after rounds of refinement, 2D classification without alignment, 3D classification with global pose search and masked 3D classification without alignment focused on the joint (**Figure S2A**). Next, particles from all datasets were extracted at 4.3 Å/px and partitioned into optics groups by hole position and grid. The initial model was separately refined against batches of particles, and batches were cleaned by 2D classification without alignment. Particles were re-extracted with re-centering at 4.3 Å/px, and the initial model was refined against all of them. Particles were then subjected to 3D classification without alignment. The best class showed residual density for additional monomers arranged in a tetrad (**Figures 3C** and **S2A**). Monomers of the tetrad were extracted at 1.45 Å/px and pooled. Refinement with global pose search, followed by duplicate removal and refinement with local pose search resulted in a map at 6.2 Å resolution. This was further classified without alignment using a mask around the arms, followed by signal subtraction of the head module, and focused refinement and 3D classification without alignment of the elbow region. After reverting to original particles, focused refinement of the head module followed by CTF refinement (beam tilt, anisotropic magnification, per-particle defocus) and Bayesian polishing were performed. A final refinement that contained the full complex inside its mask yielded a map at an overall nominal resolution of 4.6 Å.

Reconstructions of the open states and apo complex as shown in **Figure 4A** were obtained by branching off the main processing tree. Similarly, low resolution reconstructions for the apo dimer and DNA bound tetrads (**Figure 3B-3D**) were obtained by re-centering and sub-classification of classes from the main tree.

A focused reconstruction of the MukBEF–MatP–DNA head module was obtained as follows. Datasets were processed separately as shown in **Figure S2B**. Briefly, head modules from monomers within a tetrad were extracted with re-centering, subjected to multiple rounds of 3D classification, 3D refinement, CTF refinement (beam tilt, anisotropic magnification, per-particle defocus) and a final round of Bayesian polishing. Duplicate removal was performed on multiple occasions to exclude the same head in a tetrad contributing multiple times. At final stages, refinement was performed with local pose search. Using a mask around the head module, this resulted in two maps, one from pooled datasets 1 and 2 at 3.5 Å resolution, and one from dataset 3 at 3.4 Å resolution. After pooling particles, the combined refinement yielded a map at 3.3 Å resolution. Re-centering on the holocomplex with box expansion, followed by CTF refinement and Bayesian polishing yielded a final map at 3.1 Å resolution.

Maps were rendered in ChimeraX (Pettersen et al., 2021). Fourier shell correlation (FSC) for half-maps was computed in PHENIX and is shown in **Figure S2C**. Data collection and map statistics are shown in **Table S1**.

### Model building

First, a model for the head module was built into its map at 3.1 Å resolution, sharpened by B-factor compensation and FSC weighting (Rosenthal and Henderson, 2003) in RELION PostProcess. Homology models were obtained from PDB entries 3EUJ, 3EUH, 3VEA and 3IBP (Dupaigne et al., 2012; Li et al., 2010; Woo et al., 2009) using SWISS-MODEL (Waterhouse et al., 2018), and rigid body fitted using ChimeraX. PDB entry 6DFL was used as a starting model for AcpP (Kreamer et al., 2018). The hinge-proximal arm region was flexibly fitted in ISOLDE (Croll, 2018). The model was then partially rebuilt in Coot (Emsley et al., 2010), whereby the necks, joints and parts of the hinge-proximal arm were built *de novo*. The clamped DNA was modelled as poly-AT. Next, the model was subjected to phenix.real_space_refine (Afonine et al., 2018) to resolve major clashes, and annealed using ISOLDE. The model was improved by cycles of editing in ISOLDE and Coot, and automated refinement in phenix.real_space_refine using secondary structure restraints (alpha, beta, base-pair), Ramachandran restraints, without non-crystallographic symmetry (NCS) constraints. Model vs. map FSC was computed in PHENIX and is shown in **Figure S2C**.

A medium resolution model for the complete MukBEF–MatP–DNA monomer was built as follows. The head module was rigid body fitted into the 4.6 Å holocomplex map sharpened in RELION PostProcess. Homology models for hinge and elbow were obtained from PDB entries 3IBP and 6H2X (Bürmann et al., 2019; Li et al., 2010), respectively, using SWISS-MODEL. Homology models were split and flexibly fitted using ISOLDE. Connecting segments between head- and hinge-proximal arms and the elbow coiled coil were built *de novo*. There was little or no sidechain information for these segments, but their simple coiled-coil architecture and highly constrained ends allowed building of a realistic model. For example, residues of the predicted hydrophobic heptad-repeat patterns locate to the helix interfaces, and prolines locate to helix breaks. The model was improved by editing in ISOLDE and Coot, and automated refinement in phenix.real_space_refine using reference model restraints for hinge and head module, secondary structure restraints, Ramachandran restraints, and no NCS constraints. Model vs. map FSC was computed in PHENIX and is shown in **Figure S2C**.

Models for apo state and states with more open arm conformations were obtained by flexible fitting of the holocomplex reference model in ISOLDE followed by automated refinement in phenix.real_space_refine.

Low resolution models for dimers and tetrads were obtained by rigid body fitting of holocomplex monomers into multimer maps blurred to 20 Å resolution. Connecting peptides in MukF were built in Coot with a homology model of the MukF dimer based on PDB entry 3EUH (Woo et al., 2009) as a guide. The models contained some clashes at the MukF interfaces, some of which were resolved by changing sidechain rotamers without touching the main chain. Next, idealized 80 bp double-stranded DNA was generated in Coot and flexibly fitted in ISOLDE using strong distance restraints. Protein-bound regions of the DNA were replaced by the corresponding parts of the rigid-body docked medium resolution DNA models. The head bound poly-AT models were edited to match the oligonucleotide sequence. DNA was subjected to a single cycle of phenix.real_space_refine with base pair and input model restraints to resolve major geometry errors. For the tetrad with dimers directly apposed (**Figure 3D**), only two monomers were modelled due to weak density for the remaining two monomers.

Models were rendered in ChimeraX. Model statistics are listed in **Table S1**.

### *E. coli* strains

Strains are based on *E. coli* MG1655 and are listed in **Table S2**. All strains were viable in LB at 37 °C, except for *ΔmukB*, *mukB*(S1366R), *mukB*(D1407A) and mukB(E1406Q) derivatives which were cultivated at 22 °C, their permissive temperature. The strain containing cap, neck, and head cysteines for circularization of the clamp compartment (SFB202) was viable at 37 °C, but grew with a reduced rate, whereas all other strains with functional alleles grew with rates similar to WT. Pre-cultures for all experiments were grown side-by-side to stationary phase and stored at 4 °C for up to two weeks.

### Genome engineering for strain construction

For construction of acceptor strains we integrated a *pheS(T251, A294G)-hyg^R^* double selection cassette either downstream of the *mukFEB* operon (SFB047, SFB180) or replacing the *mukFEB* coding region (SFB053) by *λ*-Red recombineering (Yu et al., 2000). *pheS(T251A, A294G)* is a robust negative selection marker that encodes a mutant phenylalanyl-tRNA synthetase which confers toxicity in presence of 4-chloro-phenylalanine (4-CP) (Miyazaki, 2015). The locus was then targeted using a REXER-based strategy to introduce a C-terminal HaloTag on *mukB* and desired point mutations (Wang et al., 2016 and in preparation). The modified locus was linked to a *neo^R^* cassette downstream of the *mukFEB* transcriptional terminator. Recombinants were selected on media containing 12.5 μg/mL kanamycin and 2.5 mM 4-CP.

A marker-free *ΔmatP* allele was constructed by replacing *matP* with *pheS(T251, A294G)-hyg^R^* and subsequent cassette ejection with a double-stranded oligonucleotide coding for a *matP* in-frame deletion.

All strains were single colony purified, and verified by marker analysis, phenotype, PCR, and Sanger sequencing where appropriate.

### In vivo cross-linking

Cells were grown to exponential phase in LB (OD 0.2-0.3) if not indicated otherwise. Cultures were mixed with 30 % (w/v) ice, harvested by centrifugation, and kept cold for the duration of the experiment. 0.5 OD units of cells were washed in 500 μL of ice-cold PBS and resuspended in 50 μL PBS. Next, 1.25 μL of BMOE (20 mM stock in DMSO) were added and the suspension was incubated for 10 min on ice. The reaction was quenched by addition of 1 μL of 2-mercaptoethanol (2-ME, 1.4 M stock in water). Cells were resuspended in 50 μL of B-PER (Thermo Fisher Scientific) containing 1 mM EDTA (pH 7.4 at RT), 14 mM 2-ME, protease inhibitor cocktail (Roche), 1 μM HaloTag-TMR substrate (Promega), 0.25 U/μL Benzonase (Merck), and 0.1 U/μL ReadyLyse Lysozyme (Lucigen). The suspension was incubated for 5 min at RT and then for 10 min at 37 °C, after which 16.6 μL of 4x LDS sample buffer containing 6 % (v/v) 2-ME were added and the sample was incubated for 5 min at 95 °C. 10 μL of sample were resolved on NuPAGE 4-12 % Bis-Tris gels (Thermo Fisher Scientific) using MOPS running buffer. Gels were scanned on a Typhoon FLA9000 (GE Healthcare) with Cy3 setup. Quantification of bands was performed with Wolfram Mathematica using moving median estimation of background signal. Credible intervals were estimated from posterior distributions using a normally distributed likelihood with mean μ and standard deviation σ, a uniform prior over [0, 1] for μ and a 1/σ^2^ prior for σ.

### Chromosome entrapment assay

The *E. coli* chromosome entrapment assay was based on protocols developed for *B. subtilis* (Vazquez Nunez et al., 2019; Wilhelm et al., 2015). Stationary phase culture were inoculated into 100 mL LB and grown to OD 0.2-0.3 at 22 °C, which is the permissive temperature for ATPase deficient *mukFEB* strains. We obtained similar results at 37 °C for WT ATPase strains. Cells were harvested by centrifugation, washed in 1 mL of ice-cold PBS, and 14 OD units were resuspended in 720 μL PBS. 18 μL BMOE (20 mM in DMSO) were added, and the suspension was incubated for 10 min on ice. The reaction was quenched by addition of 14.4 μL 2-ME (1.4 M in water). Cells were resuspended in 200 μL PBS containing 10 mM EDTA (pH 7.4 at RT), protease inhibitor cocktail (Roche), and 5 μM HaloTag-TMR substrate (Promega) and incubated for 15 min at 37 °C with shaking. Samples were protected from light from now on. For input samples, 0.5 OD units from the labelling mix were collected, resuspended in 50 μL B-PER (Thermo Fisher Scientific) with 1 mM EDTA (pH 7.4 at RT), protease inhibitor cocktail, 0.25 U/µL Benzonase (Merck), and 10 U/μL ReadyLyse lysozyme (Epicentre) and incubated for 1 h at RT. Input samples were stored at -20 °C after addition of 16.6 μL 4x LDS sample buffer containing 6 % (v/v) 2-ME. Two agarose plugs per sample were formed each by mixing 100 μL cells with 100 μL low-melt agarose (2 % (w/v), freshly melted at 80 °C and equilibrated to 45 °C) in the bottom of a 2 mL tube, and incubation of the solution for 5 min on ice. Both plugs were pooled into 3 mL B-PER with 10 mM EDTA (pH 7.4 at RT), protease inhibitor cocktail, and 10 U/μL ReadyLyse lysozyme and incubated for 2.5 h at RT on a roller. Plugs were transferred into 50 mL TGES (25 mM Tris, 192 mM glycine, 10 mM EDTA (pH 7.4 at RT), 0.1 % SDS) and incubated for 2 h at RT on a roller. Plugs were mounted in a 1 % (w/v) agarose gel in TGES and subjected to electrophoresis in TGES at 10 mA cm^-2^ for 1.5 h in a chamber cooled on ice. Next, plugs were transferred into 50 mL PBS, incubated for 2 h at RT on a roller, before transfer to 2 mL tubes and melting at 80 °C for 3 min with occasional vortexing. The solution was incubated at 45 °C for 5 min, thoroughly mixed with 200 μL PBS containing 20 mM MgCl_2_ and 0.25 U/μL Benzonase and solidified on ice. Samples were incubated at 37 °C for 30 min and stored at -80 °C overnight. Next, samples were thawed at RT for 20 min and spun at 21,000 x g and 4 °C for 15 min. Extract from both plugs was combined and passed through a 0.45 μL spin filter (CoStar) by centrifugation at 10,000 x g for 1 min. Samples were brought to 1 mL with water before addition of 6 μL BSA (1 mg/mL) and 110 μL of 100 % (w/v) TCA. Tubes were incubated on ice for 30 min and precipitated protein was collected by centrifugation at 21,000 x g and 4 °C for 15 min, careful removal of the supernatant, and a second spin for 3 min to remove remaining liquid. The precipitate, which formed a haze on the tube wall, was dissolved in 20 μL 2x LDS sample buffer containing 3 % (v/v) 2-ME. Samples were incubated for 5 min at 95 °C, and 5 μL of input and 10 μL of eluate were separated on NuPAGE 3-8 % Tris-Acetate gels (Thermo Fisher Scientific) run at 4 °C and 35 mA/gel for 1.5 h. Gels were scanned on a Typhoon FLA9000 (GE Healthcare) with Cy3 setup.

Quantification of bands was performed with Wolfram Mathematica using moving median estimation of background signal. Credible intervals were estimated from posterior distributions using a normally distributed likelihood with mean μ and standard deviation σ, a uniform prior over [-10 μ, 10 μ] for μ and a 1/σ^2^ prior for σ.

## Acknowledgements

We thank J. Prince, G. Fisher and D. Sherratt (Oxford, UK) for discussions and sharing of unpublished results; R. Vazquez Nuñez and S. Gruber (Lausanne, Switzerland) for discussions and advice on the chromosome entrapment assay; all members of the Löwe and K. Nasmyth (Oxford, UK) groups for discussions; G. Cannone and all members of the LMB electron microscopy facility for excellent EM training and support; T. Darling and J. Grimmett (LMB scientific computing) for computing support. F.B. was supported by an EMBO Advanced fellowship (ALTF 605-2019).

## Author contributions

F.B. performed protein purification, cryo-EM sample preparation, cryo-EM data acquisition and analysis, model building, strain construction and all other experiments; L.F.H.F. designed the genome mutagenesis strategy; J.W.C. supervised technology development; J.L. supervised the overall study; F.B. prepared the manuscript with contributions from all authors.

## Figures

**Figure S1.**
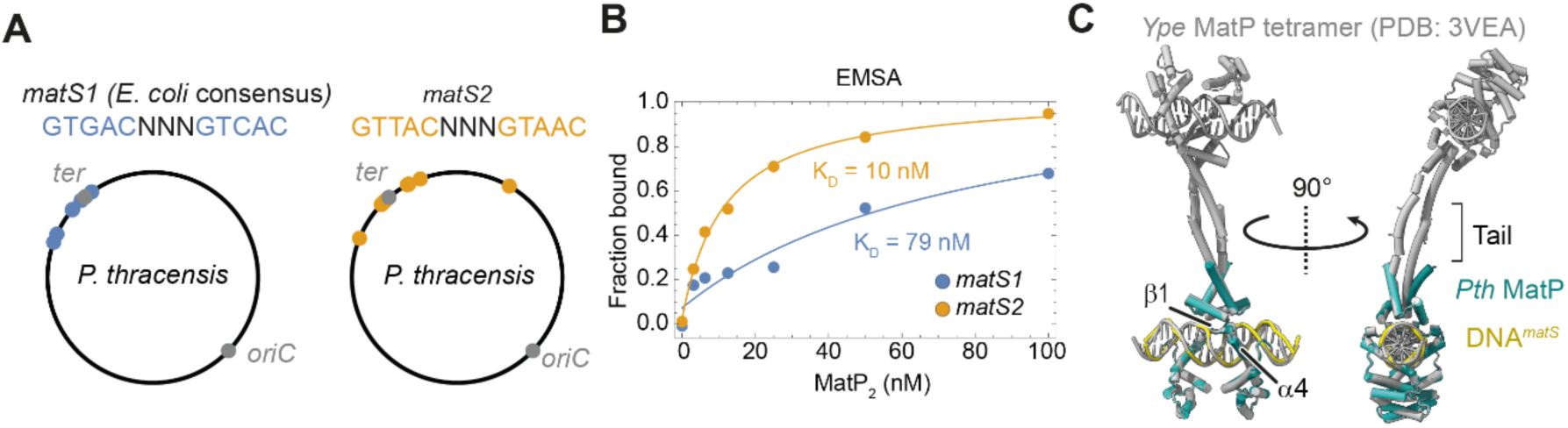
*matS* binding by *P. thracensis* MatP. Related to Figure 1. (**A**) Location of the *E. coli matS* consensus sequence (left) and the *matS* sequence used for structure determination (right) mapped onto the *P. thracensis* chromosome. (**B**) Affinities of *P. thracensis* MatP for *matS* sites shown in A as determined by EMSA. (**C**) Superimposition of MatP–*matS* in the MukBEF-bound form (colored) and a crystal structure in the absence of MukBEF (gray, PDB: 3VEA). Positions of the *matS* binding elements *α*1 and *β*1 and the C-terminal tetramerization tail are indicated. *Ype*, *Yersinia pestis*; *Pth*, *P. thracensis*.

**Figure S2.**
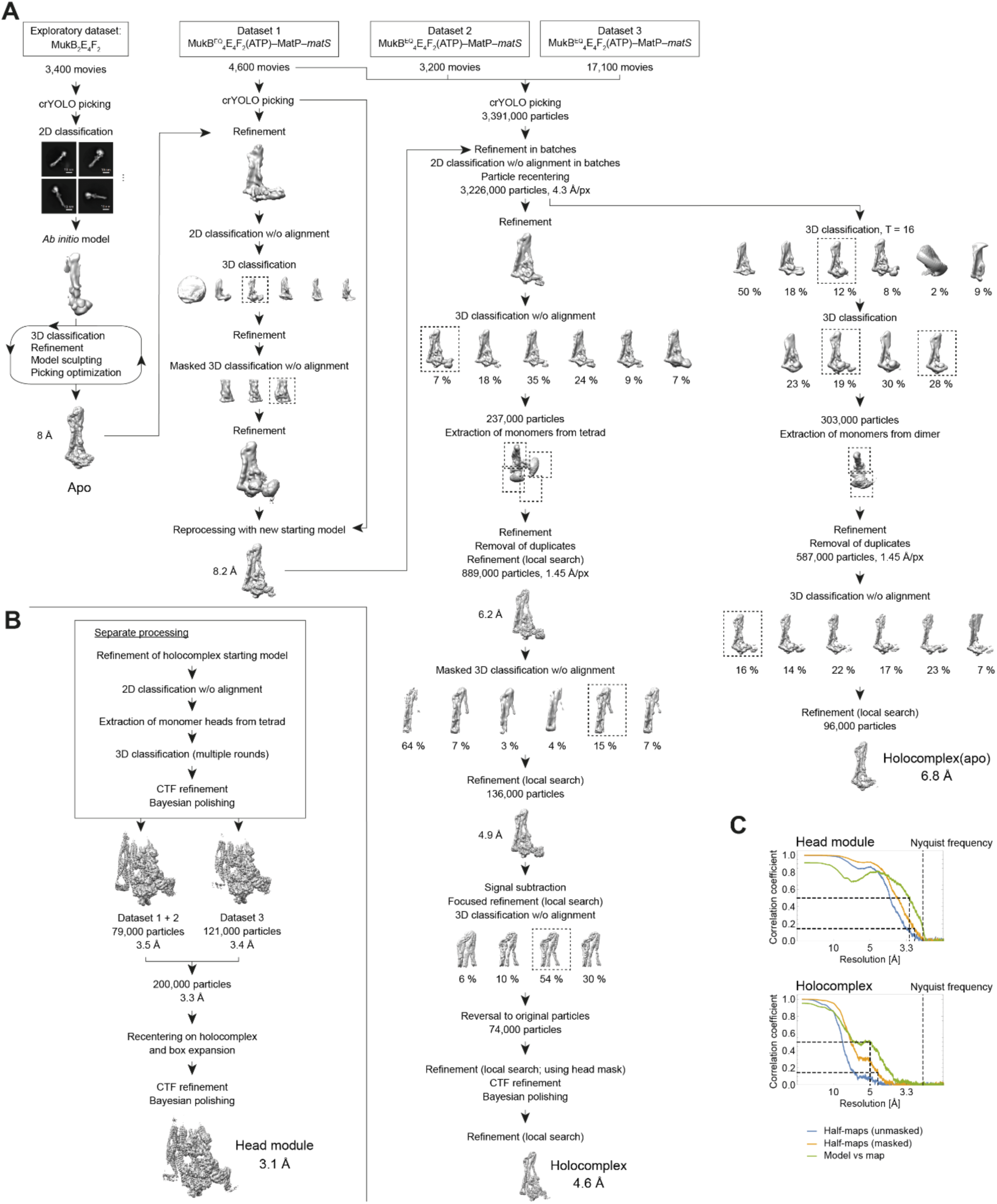
Cryo-EM data analysis workflow. Related to Figures 1-3. (**A**) The processing tree for structure determination of MukBEF–MatP–DNA and apo-MukBEF monomers. (**B**) Data processing tree for focused structure determination of the head module. (**C**) FSC curves for MukBEF–MatP–DNA head module and holocomplex structures.

**Figure S3.**
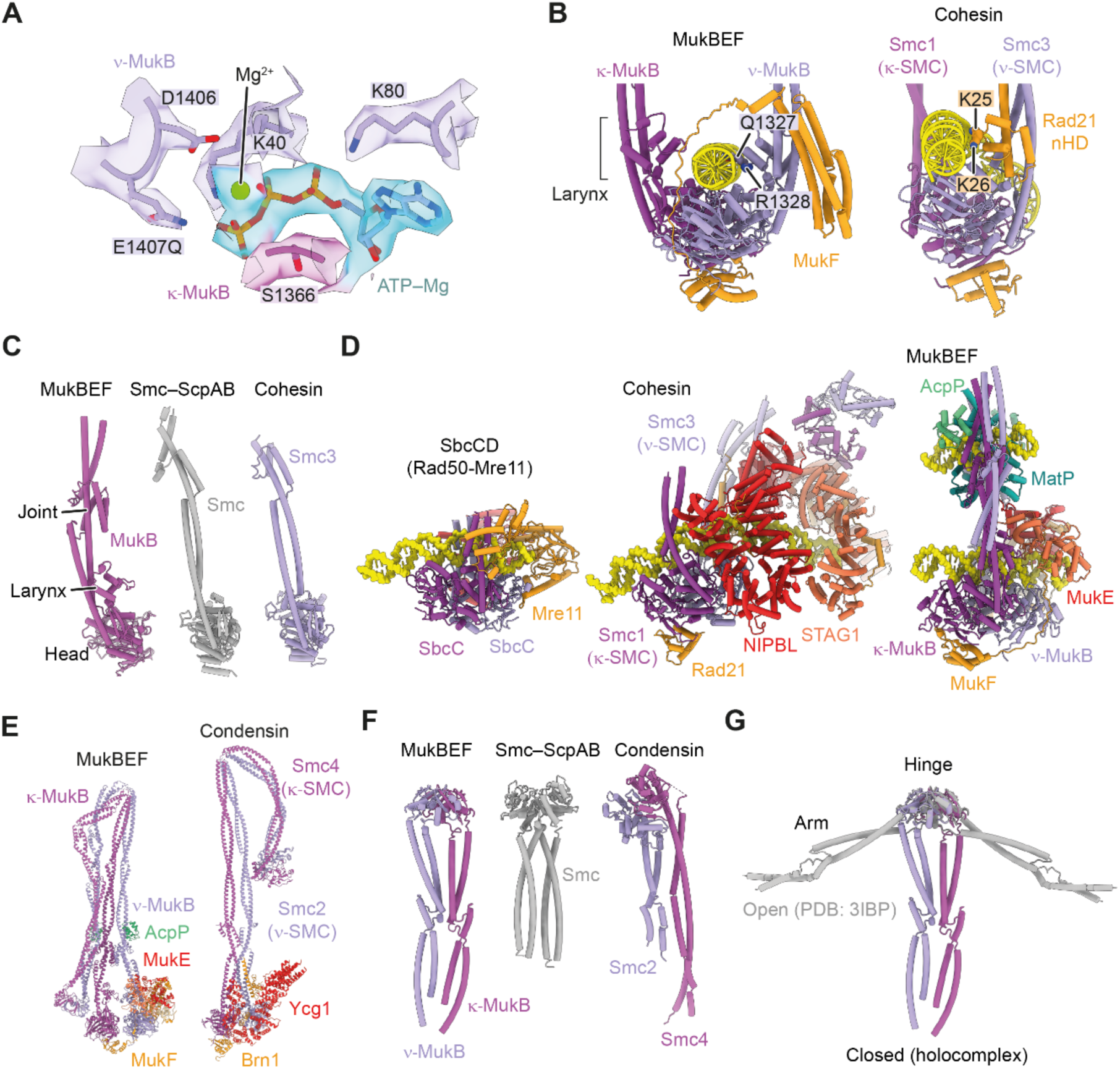
Conserved structural features and comparison with other SMC complexes. Related to Figures 1-4. (**A**) Cryo-EM density and atomic model for the nucleotide binding site at the *ν*-MukB Walker A and B motives and the *κ*-MukB signature motive. (**B**) Architecture of the neck gate in MukBEF (left) and cohesin (right, PDB: 6WG3). An asymmetric DNA contact at the larynx of MukBEF and the N-terminal helical domain of Rad21 is indicated. (**C**) Comparison of the head-proximal regions of MukB, Smc (PDB: 5XEI) and Smc3 (PDB: 6WGE). (**D**) Architecture of SMC–DNA clamps. Models were aligned on the *κ*-SMC ATPase domain. PDB: 6S85, 6WG3. (**E**) Comparison of apo-MukBEF and apo-condensin (PDB: 6YVU). (**F**) Comparison of the hinge-proximal regions of MukB, Smc (PDB: 4RSJ) and Smc2/4 (PDB: 4RSI). (**G**) Superimposition of the hinge-proximal region of MukB in an open conformation (PDB: 3IBP) and in the closed conformation.

**Figure S4.**
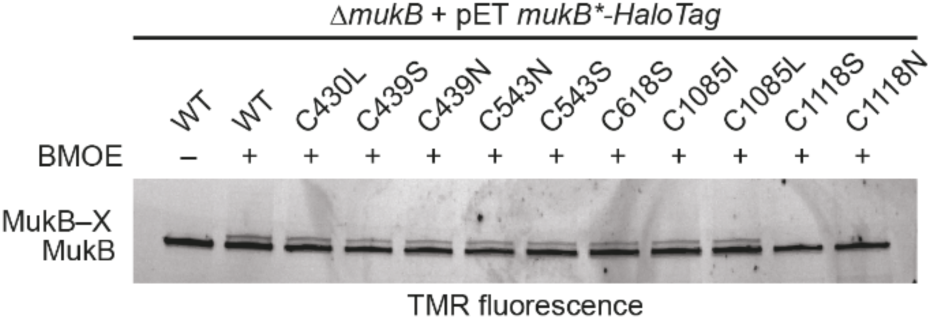
Background cross-linking of endogenous MukB cysteines. Related to Figure 5. Screening of endogenous cysteines for background cross-linking. *E. coli ΔmukB* was transformed with plasmids carrying *mukB-HaloTag* variants under control of a T7 promoter. Leaky expression in the strain lacking a T7 RNA polymerase gene produced about 40 % MukB-HaloTag compared to *mukB-HaloTag* expressed from the endogenous locus. The *mukB* null phenotype was complemented in all cases. Cells were treated with BMOE and proteins were detected by in-gel fluorescence. C1118S and C1118N abolished background cross-linking with a low-molecular weight protein.

**Figure S5.**
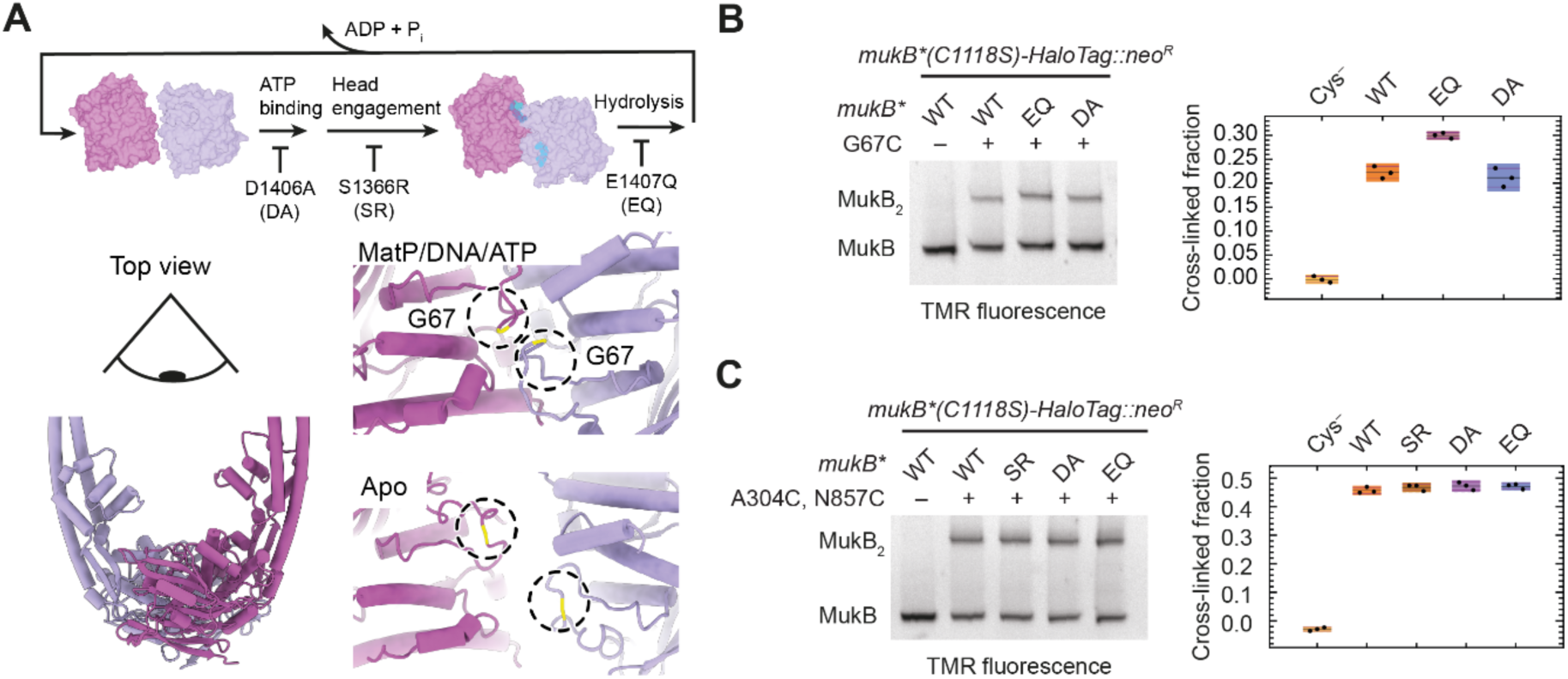
*In vivo* head engagement and arm folding in ATPase mutants. Related to Figures 5 and 6. (**A**) Schematic of the ATP hydrolysis cycle and blocking mutations (top). Location of the head engagement sensor residue G67 in MatP/DNA/ATP and apo states (bottom). S1366R (‘SR’) blocks head engagement, D1406A (‘DA’) blocks ATP binding, E1407Q (‘EQ’) blocks ATP hydrolysis. (**B**) BMOE cross-linking of strains carrying the head engagement sensor mutation G67C and ATPase blocking mutations. Cells were grown in LB for 1.5 h at 37 °C before cross-linking. In-gel fluorescence is shown on the left, and quantification of three technical replicates is shown on the right. Black lines indicate means, purple lines indicate standard deviations, and colored bars indicate 95 % credible intervals. (**C**) BMOE cross-linking of arm folding sensor strains carrying ATPase mutations. As in B.

**Figure S6.**
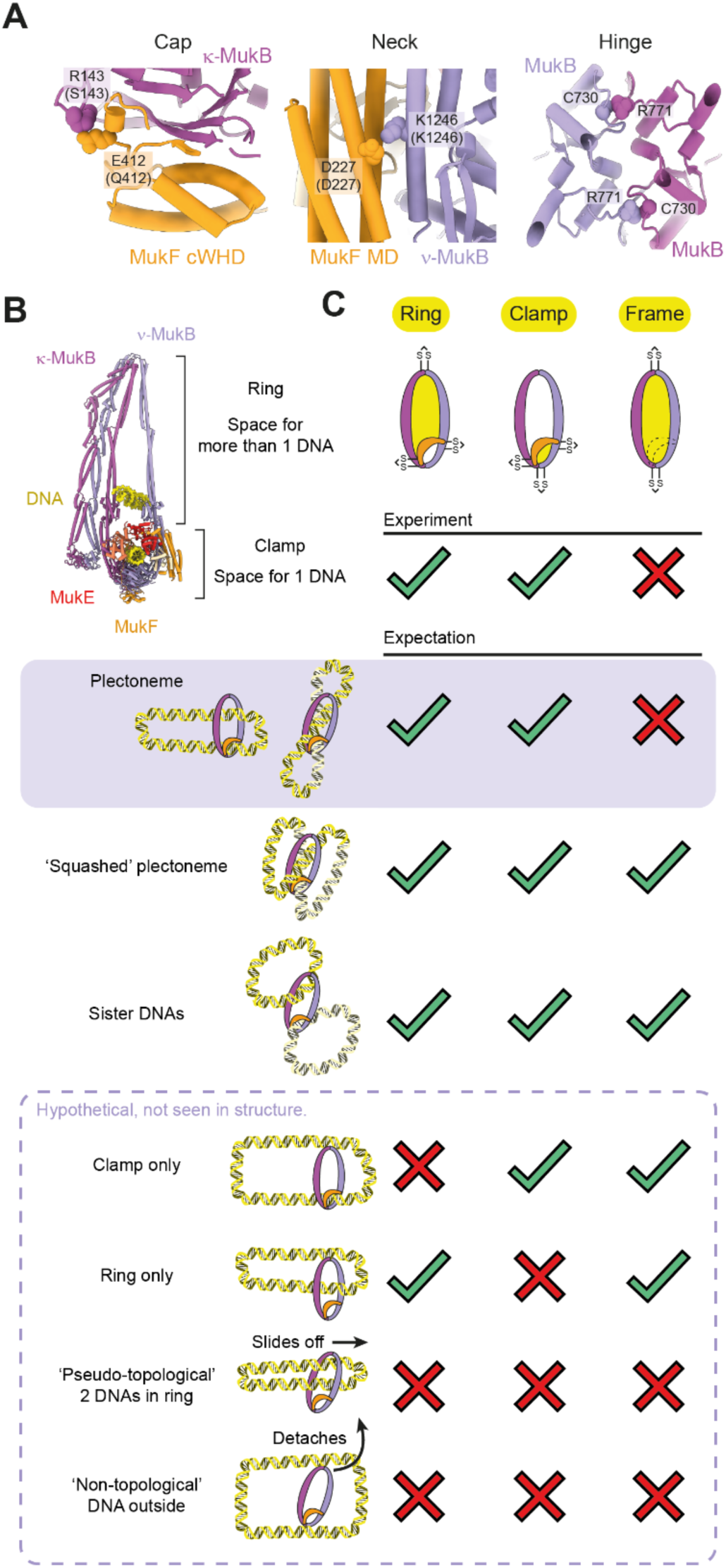
Chromosome entrapment by MukBEF. Related to Figure 6. (**A**) Design of cap, neck and hinge cross-links. Location of residues chosen for cysteine mutagenesis at the hinge (left, PDB: 3IBP), cap (middle), and neck (right) is shown. Labels for cap and neck are shown for the *P. thracensis* structure and corresponding *E. coli* residues are in parentheses. (**B**) Space for hypothetical accommodation of additional DNA double strands. Only the ring compartment is large enough to embrace more than one DNA. (**C**) Comparison of experimentally observed and expected catenanes for different DNA binding topologies. The topology consistent with the experimental data is highlighted. For clarity, the double-locked plectoneme topology is also shown as a simplified version without DNA crossings. Hypothetical ‘pseudo-topological’ and ‘non-topological’ DNA loops do not produce catenanes. Their tentative formation in addition to catenated forms is therefore not excluded by the data.

## Tables

**Table S1.** Cryo-EM data collection and model statistics.

|  | Head module | Holocomplex | Holocomplex (apo) | Holocomplex (partially open) | Holocomplex (open) | Tetrad | Tetrad (apposed) | Dimer (apo) |
| --- | --- | --- | --- | --- | --- | --- | --- | --- |
|  | EMD-12656<br>PDB 7NYW | EMD-12657<br>PDB 7NYX | EMD-12658<br>PDB 7NYY | EMD-12658<br>PDB 7NYZ | EMD-12658<br>PDB 7NZ0 | EMD-12662<br>PDB 7NZ2 | EMD-12663<br>PDB 7NZ3 | EMD-12664<br>PDB 7NZ4 |
| **Data collection and processing** |  |  |  |  |  |  |  |  |
| Magnification | 81,000 |  |  |  |  |  |  |  |
| Voltage (kV) | 300 |  |  |  |  |  |  |  |
| Electron fluence (e-/Å²) | 40 |  |  |  |  |  |  |  |
| Defocus range (µm) | -1 to -3 |  |  |  |  |  |  |  |
| Pixel size (Å) | 1.07 |  |  |  |  |  |  |  |
| Symmetry imposed | C1 |  |  |  |  |  |  |  |
| Initial particle images (no.) | 3,391,688 |  |  |  |  |  |  |  |
| Final particle images (no.) | 200,438 | 74,064 | 96,150 | 41,109 | 60,245 | 12,010 | 8,561 | 4,197 |
| Map resolution (Å) | 3.1 | 4.6 | 6.8 | 6.5 | 6.3 | 11 | 11 | 13 |
| FSC threshold | 0.143 | 0.143 | 0.143 | 0.143 | 0.143 | 0.143 | 0.143 | 0.143 |
| **Model** |  |  |  |  |  |  |  |  |
| Initial model used (PDB code) | 3EUJ, 3EUH, 3VEA, 3IBP, 6DFL | Head module, 6H2X | Holocomplex | Holocomplex | Holocomplex | Holocomplex | Holocomplex | Holocomplex (apo) |
| Model resolution (Å) | 3.25 | 5.0 | 7.5 | 8.1 | 7.3 | — | — | — |
| FSC threshold | 0.5 | 0.5 | 0.5 | 0.5 | 0.5 | — | — | — |
| Map sharpening B factor (Å²) | -33 | -87 | -174 | -162 | -157 | — | — | — |
| **Model composition** |  |  |  |  |  |  |  |  |
| Non-hydrogen atoms | 24,792 | 36,100 | 31,563 | 36,100 | 36,100 | 148,563 | 74,338 | 63,192 |
| Protein residues | 2,795 | 4,186 | 3,910 | 4,186 | 4,186 | 16,752 | 8372 | 7824 |
| Nucleic acid residues | 104 | 104 | — | 104 | 104 | 614 | 312 | — |
| Ligands | PNS: 2<br>ATP: 2<br>Mg: 2 | PNS: 2<br>ATP: 2<br>Mg: 2 | PNS: 2 | PNS: 2<br>ATP: 2<br>Mg: 2 | PNS: 2<br>ATP: 2<br>Mg: 2 | PNS: 8<br>ATP: 8<br>Mg: 8 | PNS: 4<br>ATP: 4<br>Mg: 4 | PNS: 4 |
| **R.m.s. deviations** |  |  |  |  |  |  |  |  |
| Bond lengths (Å) | 0.004 | 0.004 | 0.004 | 0.009 | 0.005 | 0.004 | 0.004 | 0.005 |
| Bond angles (°) | 0.867 | 0.926 | 0.942 | 1.102 | 0.969 | 0.929 | 0.925 | 0.954 |
| **Validation** |  |  |  |  |  |  |  |  |
| MolProbity score | 1.57 | 1.78 | 1.99 | 1.77 | 1.82 | 1.85 | 1.83 | 2.01 |
| Clashscore | 5.9 | 10.9 | 14.51 | 10.63 | 12.31 | 12.96 | 12.37 | 15.24 |
| Poor rotamers (%) | 0 | 0 | 0 | 0.08 | 0.03 | 0.04 | 0 | 0.03 |
| **Ramachandran plot** |  |  |  |  |  |  |  |  |
| Favored (%) | 96.31 | 96.57 | 95.35 | 96.54 | 96.64 | 96.55 | 96.58 | 95.31 |
| Allowed (%) | 3.69 | 3.43 | 4.65 | 3.46 | 3.36 | 3.45 | 3.42 | 4.68 |
| Disallowed (%) | 0 | 0 | 0 | 0 | 0 | 0 | 0 | 0.01 |

**Table S2.**
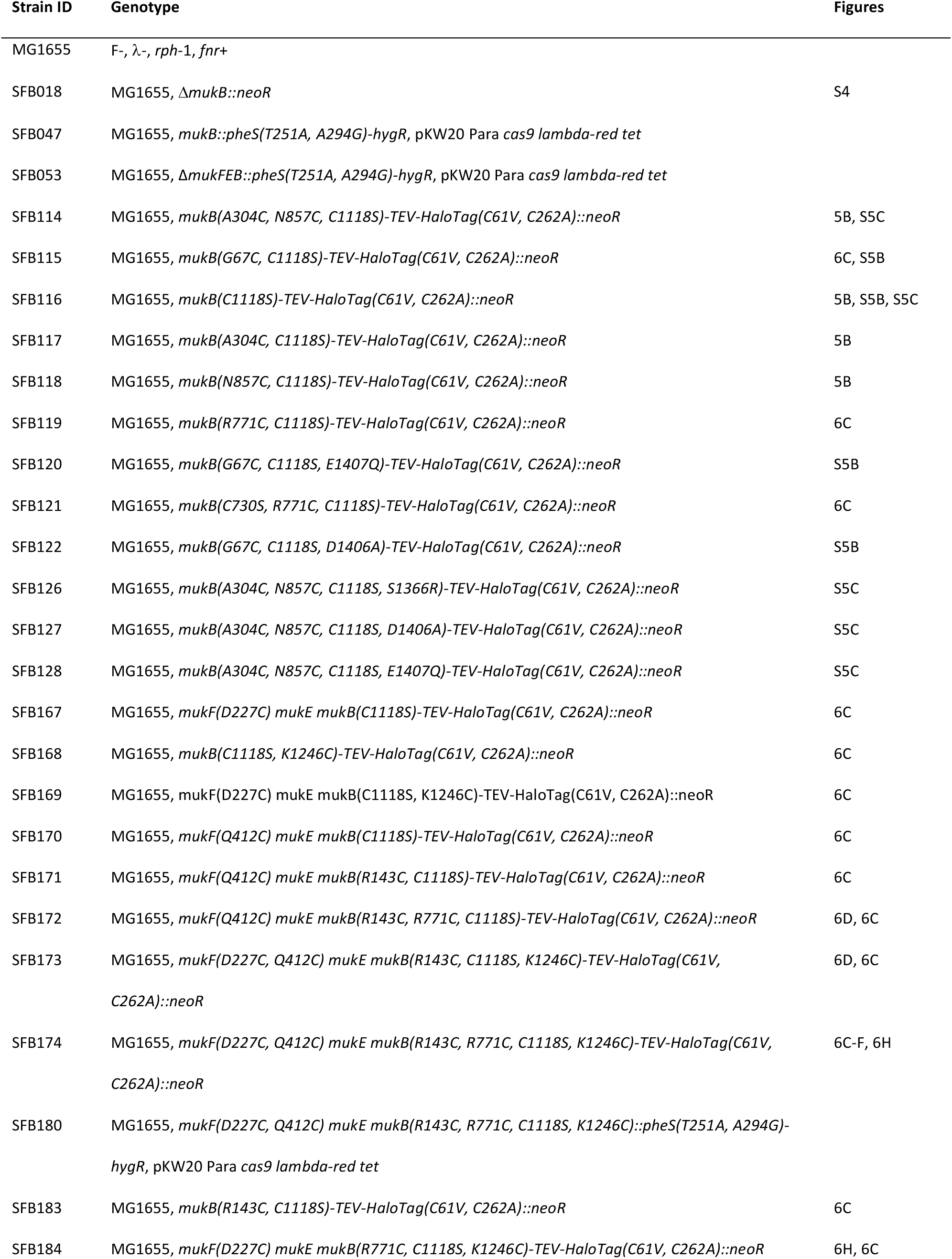
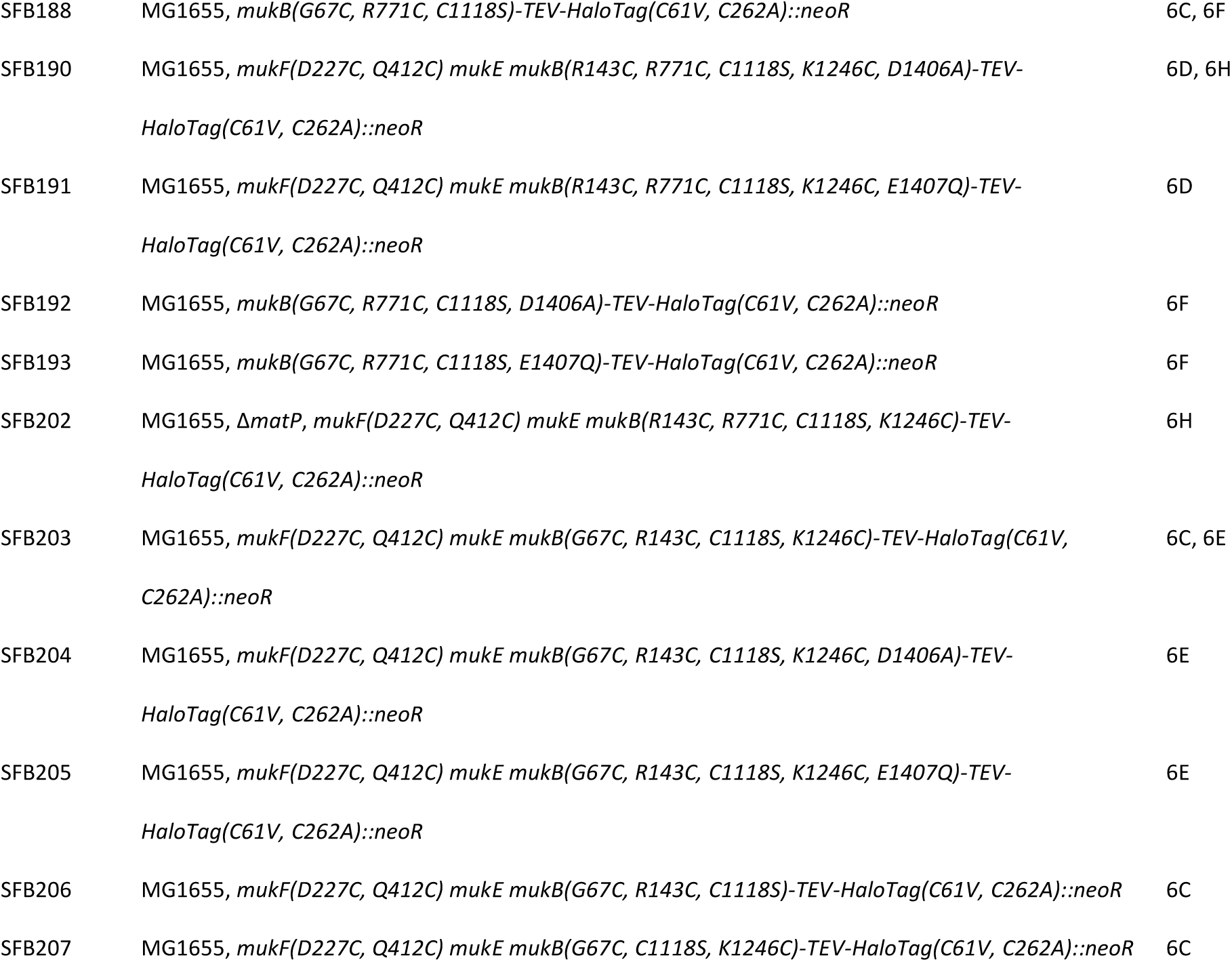
Bacterial strains.

